# RET Receptor Tyrosine Kinase Promotes Breast Cancer Metastasis to the Brain and RET Inhibitors Pralsetinib and Selpercatinib Suppress Breast Cancer Brain Metastases

**DOI:** 10.1101/2025.10.07.680986

**Authors:** Angelina T. Regua, Shivani Bindal, Mariana Najjar, Phi-Long Tran, Joshua Cha, Syed Shahmeer Shams, Hui-Wen Lo

**Affiliations:** Vivian L. Smith Department of Neurosurgery, McGovern Medical School, The University of Texas Health Science Center at Houston, Houston, Texas, United States; The University of Texas MD Anderson Cancer Center UTHealth Houston Graduate School of Biomedical Sciences, Houston, Texas, United States; Department of Cell Biology and Genetics, Naresh K. Vashisht College of Medicine, Texas A&M University Health Science Center, College Station, Texas, United States; Department of Translational Medicine, Naresh K. Vashisht College of Medicine, Texas A&M University Health Science Center, College Station, Texas, United States; Institute of Biosciences & Technology, Texas A&M University Health Science Center, Houston, Texas, United States

## Abstract

Patients with breast cancer brain metastases (BCBM) exhibit dismal prognosis, largely due to the insufficient biological understanding of BCBM and the scarcity of therapeutics that can penetrate the blood-brain barrier. This study was focused on Rearranged during transfection (RET) receptor tyrosine kinase that has been implicated in tumorigenesis and metastatic progression of several solid tumor types including, non-small cell lung cancer (NSCLC), thyroid carcinomas, and luminal breast cancer subtypes. FDA-approved selective RET inhibitors, pralsetinib and selpercatinib, have demonstrated potent intracranial activity in brain metastases from NSCLC and thyroid cancer; however, their efficacy in BCBM has not been investigated. Here, we report that RET activation is elevated in brain metastases of breast cancer patients compared to matched primary tumors (N=30), and in three brain-tropic breast cancer cell lines compared to the parental lines. High RET pathway activation is associated with shorter overall metastasis-free survival and brain metastasis-free survival in patients with HER2-enriched and triple-negative breast cancer (TNBC). Using isogenic TNBC cells lines RET overexpression, we demonstrated that RET strongly promotes their preferential metastasis to the brain in mice with intracardiac injections to tumors cells. Using intracranial tumor implantation of the isogenic lines, we further found that RET significantly enhances the formation and progression of brain tumors *in vivo*. Moreover, we report that selective RET inhibition using pralsetinib and selpercatinib significantly reduces cell viability, enhances apoptosis, and attenuates migration of brain-tropic breast cancer cells *in vitro*. Using two mouse studies that model multi-organ metastases and breast tumor formation in the brain, we observed that RET inhibition significantly prevented the circulating tumor cells from forming brain metastases and suppressed the growth of intracranially implanted tumor cells, but did not significantly inhibit the progression of well-established brain metastases. Together, our findings demonstrated that RET is highly activated in BCBM and functioning as a novel mediator of BCBM, and that RET plays a new role as a viable therapeutic target for BCBM.

## INTRODUCTION

Breast cancer is the most commonly diagnosed cancer and the second-leading cause of cancer related deaths among American women [1, 2]. Among the various subtypes, HER2-enriched and triple-negative breast cancer (TNBC) subtypes are the most aggressive, exhibiting the marked propensity for distant metastases, particularly to the brain [3, 4]. Breast cancer brain metastases (BCBM) occur in approximately 10-30% of metastatic breast cancer cases and are associated with dismal prognoses, with a median survival time of 7.9 months, which is significantly lower than the median survival for metastases to the lung (25 months), liver (31 months), or bone (36 months) [4, 5]. Currently, there is no recommended standard of care for BCBM, and the treatment landscape remains severely limited due to the lack of actionable molecular targets and the challenge of delivering therapeutics across the blood-brain and blood-tumor barriers (BBB/BTB) [6, 7]. These limitations underscore the urgent need to identify novel therapeutic candidates in BCBM that are actionable with brain-penetrant compounds.

Rearranged during transfection (RET) is a receptor tyrosine kinase encoded by the RET gene and is prone to forming oncogenic RET fusions in various solid cancer types [8]. RET is activated upon binding of its cognate ligand, glial cell line-derived neurotrophic factor (GDNF), to the co-receptor GDNF family receptor alpha-1 (GFR1α), which facilitates the dimerization of RET monomers through enhanced proximity. This induces cross-phosphorylation and activation of the RET intracellular tyrosine kinase domains, thereby propagating downstream signaling that promote cell proliferation and survival. Although RET expression is upregulated and essential during embryonic development, it is downregulated in most adult tissues [8]. Aberrant RET expression or activation has been implicated in tumorigenesis and metastatic progression in several solid tumor types, including non-small cell lung cancers (NSCLC), thyroid cancers and luminal breast cancers, characterizing RET as an important therapeutic target [8].

Due to the prevalence of RET aberrations in multiple metastatic cancers, significant efforts have been made to develop selective RET inhibitors. Multikinase inhibitors (MKI) demonstrated activity against RET but were associated with substantial off-target toxicities, warranting the development of selective RET inhibitors [8]. Two orally bioavailable, selective RET small molecule inhibitors (RETis), pralsetinib and selpercatinib, have since received FDA approval for RET-altered NSCLC and thyroid carcinomas. Pralsetinib (Gavreto®) is FDA-approved for adults with RET-altered NSCLC, as well as for patients with advanced or metastatic RET-altered thyroid cancers who require systemic therapy and are refractory to radioactive iodine [9–13]. The clinical development of pralsetinib is ongoing, with Phase I/II trials in NSCLC and medullary thyroid carcinomas (NCT03037385) and a Phase III trial initiated in 2020 (NCT04222972) for patients with advanced/metastatic NSCLC. Selpercatinib (Retevmo®) is approved for adults with RET-altered NSCLC, and for adult and pediatric patients with RET-altered metastatic medullary thyroid cancers, and RET-altered metastatic thyroid cancers requiring systemic therapy and are refractory to radioactive iodine [14]. Notably, selpercatinib became the first RETi to receive tumor-agnostic FDA approval for adults with advanced or metastatic solid tumors harboring RET fusions [15]. Importantly, both pralsetinib and selpercatinib have been reported to penetrate the BBB/BTB, as supported by clinical trial data demonstrating potent intracranial activity in patients with NSCLC and thyroid cancer [16–19]. However, the therapeutic utility of selective RETi in BCBM, particularly in HER2-enriched and triple-negative subtypes, remains to be elucidated.

In this present study, we identify a positive correlation between elevated RET activity and BCBM, as well as an association between high RET pathway activation and shorter overall and brain metastasis-free survival in aggressive breast cancer subtypes, including HER2-enriched and TNBC. Our *in vitro* data support these findings, demonstrating significantly higher levels of active RET in BCBM and brain-tropic breast cancer cells compared to their matched primary tumors or parental lines, respectively. RET overexpression in breast cancer cells promoted preferential metastasis to the brain in the intracardiac injection metastasis mouse model and enhanced the growth of intracranially implanted breast tumors *in vivo*. Importantly, we showed that pharmacologic inhibition of RET using pralsetinib or selpercatinib reduced cell viability and induced apoptosis in brain-tropic breast cancer cell lines. These inhibitors also significantly impaired cell migration, though no significant effect on invasion was observed *in vitro*. Using two mouse studies that model multi-organ metastases and breast tumor formation in the brain, we observed that RET inhibition significantly prevented the circulating tumor cells from forming brain metastases and suppressed the growth of intracranially implanted tumor cells, but did not significantly inhibit the progression of well-established brain metastases, suggesting a need for RET-based combination therapies.

## MATERIALS AND METHODS

### Cell lines and cell culture

Brain-tropic SKBR3 (SKBRM) cells were provided by Drs. Fei Xing and Kounosuke Watabe at Wake Forest School of Medicine [20]. Parental CN34 and MDA-MB-231, as well as their respective brain-tropic sublines (CN34-BrM, MDA-MB-231-BrM) were provided by Dr. Joan Massagué at MIT [21]. All other cell lines were purchased from American Type Culture Collection (ATCC, Manassas, VA), maintained in appropriate growth media, and validated using standard methods. Cell lines were routinely checked for mycoplasma using a mycoplasma detection kit (InvivoGen; San Diego, CA).

### Pearson Correlation, Metastasis-Free Survival, and Statistical Analyses

Publicly available breast tumor RNA-Seq expression profiles were retrieved from the Gene Expression Omnibus (GEO) (GSE2034, GSE2603, GSE5327, GSE12276, GSE14020) with metastasis-free survival information. Median centering was used to generate a RET Activation Signature comprised of known genes regulated by RET pathway activity (R-HSA-8853659.2). A Breast-to-Brain-Metastasis Signature, comprised of genes altered in brain-tropic breast cancer cells, was also used [21]. For comparison of RET activation in patients without metastases and with brain metastases, RET activation scores were calculated, and patients were sorted based on brain metastasis status before the activation scores were plotted on GraphPad Prism 10. For correlation analyses, the median-centered scores for Breast-to-Brain-Metastasis Signature and RET Activation Signature were plotted for each patient and Pearson correlation was used to calculate the coefficient of determination (R2). Regression analysis was performed to calculate p-values. For Kaplan-Meier survival analyses, RET activation scores were calculated, and patients were stratified based on high or low RET activation (50:50) based on the median activation score and plotted using GraphPad Prism 10. The Log-rank test was used to determine statistical significance.

### Immunohistochemistry (IHC)

Mouse brains were frozen at −80⁰C using optimal cutting temperature (OCT) compound (Fisher Scientific; Waltham, MA, USA) and subjected to IHC as previously described, or matched primary breast tumors and BCBM paraffin embedded sections obtained from Dr. Wencheng Li (Wake Forest School of Medicine) [22]. Anti-p-RET (Y905) (R&D AF3269,1:10); Anti-Ki67 (CST 9027s, 1:400).

### Pharmacological Inhibition and Cell Viability Assays

RET inhibitors (pralsetinib and selpercatinib) were purchased from Adooq BioScience (Irvine, CA) and dissolved in dimethyl sulfoxide (DMSO). For cell viability assays, 2 x 10^5^ cells were seeded onto a Greiner white flat-bottom 96-well plate. Cells were treated with increasing doses of either RET inhibitor for 48 hours and subjected to Cell Titer Blue cell viability assay (ProMega; Madison, WI, USA) as per manufacturer’s protocol.

### Annexin V flow cytometry

Cells were seeded onto 10-cm dishes (1 x 10^6^ cells/dish) and allowed to adhere overnight at 37°C. Cells were treated with Vehicle, pralsetinib, or selpercatinib for 48 hours, after which floating and adherent cells were stained with Annexin V-CF Blue and PI apoptosis detection kit (Abcam, ab214485) as per manufacturer’s protocols.

### Western Blot

Cells were lysed using RIPA buffer supplemented with Halt protease and phosphatase inhibitor and nuclease (MilliporeSigma; Burlington, MA, USA). Lysates were clarified by centrifugation (>20,000 x g for 20 minutes at 4⁰C) and subjected to Bradford assay (Bio-Rad; Hercules, CA, USA) before preparing in Laemmli buffer and boiling at 95⁰C for 5 minutes. 100μg of whole cell lysate was loaded per lane using Bio-Rad pre-cast TGX gels and subjected to wet transfer with Tris-glycine transfer buffer (20% methanol). The following antibodies were used: p-RET (Y905) (CST 3221s, 1:500); p-AKT (5473) (CST 9271s, 1:1000); p-ERK (T202/Y204) (CST 91015, 1:1000); p-STAT3 (CST 9131L, 1:1000); Vinculin (CST 13901s, 1:5000); RET (CST 3220s, 1:500); RET (CST 3223s, 1:1000); AKT (CST 9272s, 1:1000); ERK (CST 4695s, 1:1000); STAT3 (CST 9139, 1:1000); β-actin (CST 3700s, 1:5000); α-tubulin (Sigma-Aldrich T6074, 1:5000–10,000). Western blots were imaged using Bio-Rad ChemiDoc imaging system. Densitometry was performed using the ChemiDoc analysis software.

### Migration and Invasion Assays

For migration assays, cells were seeded (2 – 5 ×10^5^ cells/well) onto 6-well plates and allowed to adhere overnight at 37°C. The following day, a scratch was made using a 200µL pipette tip and media was immediately replaced with drug-containing media. Wells were imaged for T0 using the Keyence BZ-X microscope and returned to the incubator for 24 hours, after which they were imaged again for T24. To account for effects of cell viability on migration, cell viability assays were performed in parallel to the migration assays. All migration data was normalized to the respective cell viability data to calculate net migration rate. Invasion assays were carried out using the Sigma Aldrich QCM ECMatrix Cell Invasion Assay (ECM555) or the Abcam Cell Invasion Assay (Basement Membrane; ab235697) as per manufacturer’s protocols. Briefly, cells were seeded on the top chamber in FBS-free growth media and treated with pralsetinib or selpercatinib. In the bottom wells, media containing FBS (10%) was used as the chemoattractant. Cells were treated overnight (∼16 hours) at 37°C before plates were assayed for invasion. As with the migration assays, cell viability assays were performed in parallel to account for effects of cell proliferation on the invasion data. All invasion data was normalized to the respective cell viability data to calculate net invasion rate.

### Generation of Cell Lines with Stable RET Overexpression

MDA-MB-231-luc cells were seeded onto a 12-well dish and infected with lentivirus for RET overexpression at MOI 2. Virus particles were generated by VectorBuilder (Chicago, IL). After transduction, cells were subjected to antibiotic selection for one week. Transduced cells were validated for RET overexpression via Western blot analysis.

### Intracardiac Inoculation Mouse Studies

All animal studies were completed with approval from the Institutional Animal Care and Use Committee. All colonies of female athymic nude mice (#002019, Jackson Laboratory) were housed in a pathogen-free facility in a 12:12-h light/dark cycle and fed irradiated rodent chow ad libitum. Anesthetized mice were inoculated with 2×10^5^ MDA-MB-231-BrM cells and subjected to bioluminescent imaging (BLI) bi-weekly as previously described [23]. For the pre-brain metastasis treatment study, mice began receiving daily oral administration of vehicle (100% polyethylene glycol-300; PEG-300) or selpercatinib (30 mg/kg) one day after inoculation. For post-brain metastasis treatment, mice were allowed to develop metastases for 10-14 days, after which they were randomized based on brain BLI and began receiving daily oral administration of vehicle or selpercatinib. All treatments were administered on a 5 days on/2 days off weekly regimen. Animal weights were measured twice/week.

### Intracranial Injection Mouse Studies

We performed intracranial injection of breast cancer cells as previously described [24]. Briefly, anesthetized female athymic nude mice were intracranially inoculated with MDA-MB-231-BrM cells (1×10^4^ cells/brain). Mice began receiving daily oral administration of vehicle (100% PEG-300) or Pralsetinib (10 mg/kg) one day after inoculation. Mice were subjected to BLI imaging twice/week. All treatments were administered on a 5 days on/2 days off weekly regimen. Animal weights were measured twice/week.

### Intracardiac and Intracranial Injection of MDA-MB-231 cells with RET overexpression

Intracardiac injection and intracranial of MDA-MB-231-RET cells were performed as described earlier. Animals were subjected to bi-weekly bioluminescent IVIS imaging to track tumor growth. Animal weights were recorded twice/week.

### Alanine Transaminase Assay

Hepatotoxicity was evaluated using the Alanine Transaminase (ALT) Colorimetric Assay Kit (Cayman Chemical; Ann Arbor, MI, USA) as previously described [22, 25]. Samples were run in technical triplicate and five samples per treatment group were assayed. Toxicity thresholds indicated were based on ALT activity level found in nude mice treated with sublethal doses of thioacetamide, a compound known to induce acute hepatocellular toxicity [26].

### TUNEL assay

Apoptotic DNA fragmentation was analyzed by using Click-iTTM Plus TUNEL assay (CAT#C10618, Invitrogen, Oregon, 97402, USA) following the manufacturer’s protocol. Briefly, 5mm-thick slices of mouse brain tumor were fixed using 4% paraformaldehyde and permeabilized with proteinase K, then washed twice with PBS. Slices were subjected to the TdT reaction, followed by the Click ITTM Plus reaction. To label nuclei, slices were stained and mounted with DAPI Flouromount-GÒ (CAT#101492-494, Southern Biotech, AL 35260, USA). Slices were then imaged by the All-in-One Fluorescence Microscope KEYENCE BZ-X800 and analyzed by using ImageJ software.

### Statistical Analysis

Data were analyzed using CompuSyn (ComboSyn, Inc.; Paramus, NJ, USA), Prism 10 (GraphPad Software; San Diego, CA, USA), and Excel 2016 (Microsoft; Redmond, WA, USA). Descriptive statistics are presented as mean ± SEM. One-way and two-way ANOVA were performed using Prism 8. All experiments were performed at least 3 times to acquire SEM.

## RESULTS

### RET activity is enriched in patients with BCBM and is associated with shorter overall and brain metastasis-free survival

To determine the prevalence and potential clinical significance of RET pathway activation in patients with BCBM, we performed data mining analyses using a curated breast cancer patient dataset compiled from publicly available Gene Expression Omnibus (GEO) datasets (GSE2034, GSE2603, GSE5327, GSE12276, GSE14020). Using a previously published RET activation gene signature, we found that patients with BCBM exhibit significantly higher RET activation compared to those without metastases (Fig. 1A), suggesting that RET pathway activity may be associated with BCBM [27, 28]. To further assess this association, we conducted a Pearson correlation analysis between the RET activation signature score and the score of a previously published Breast-to-Brain metastasis gene signature. Our analysis revealed a significant positive correlation between RET pathway activation and Breast-to-Brain Metastasis signature in breast cancer patients (Fig. 1B), suggesting a potential pro-oncogenic role for RET in promoting brain metastases among breast cancer patients. We next evaluated the prognostic relevance of RET pathway activity by performing Kaplan-Meier survival analyses on our curated breast cancer patient cohort. We found that increased RET activation is significantly associated with shorter overall metastasis-free survival (Fig. 1C) and shorter brain metastasis-free survival (Fig. 1D), underscoring its potential role as a negative prognostic marker. To corroborate these findings at the protein level in tumor tissues, we performed immunohistochemical (IHC) staining for activated RET (p-RET Y905) on 15 matched pairs of primary breast tumors and their corresponding BCBM samples (Fig. 1E). Notably, we found that 94% (14/15) of the BCBM specimens exhibited significantly increased p-RET (Y905) expression compared to their matched primary tumors (Fig. 1F), further supporting the enrichment of RET signaling in brain metastases. Collectively, these findings suggest that elevated RET activity may contribute to the development of BCBM.

**Figure 1:**
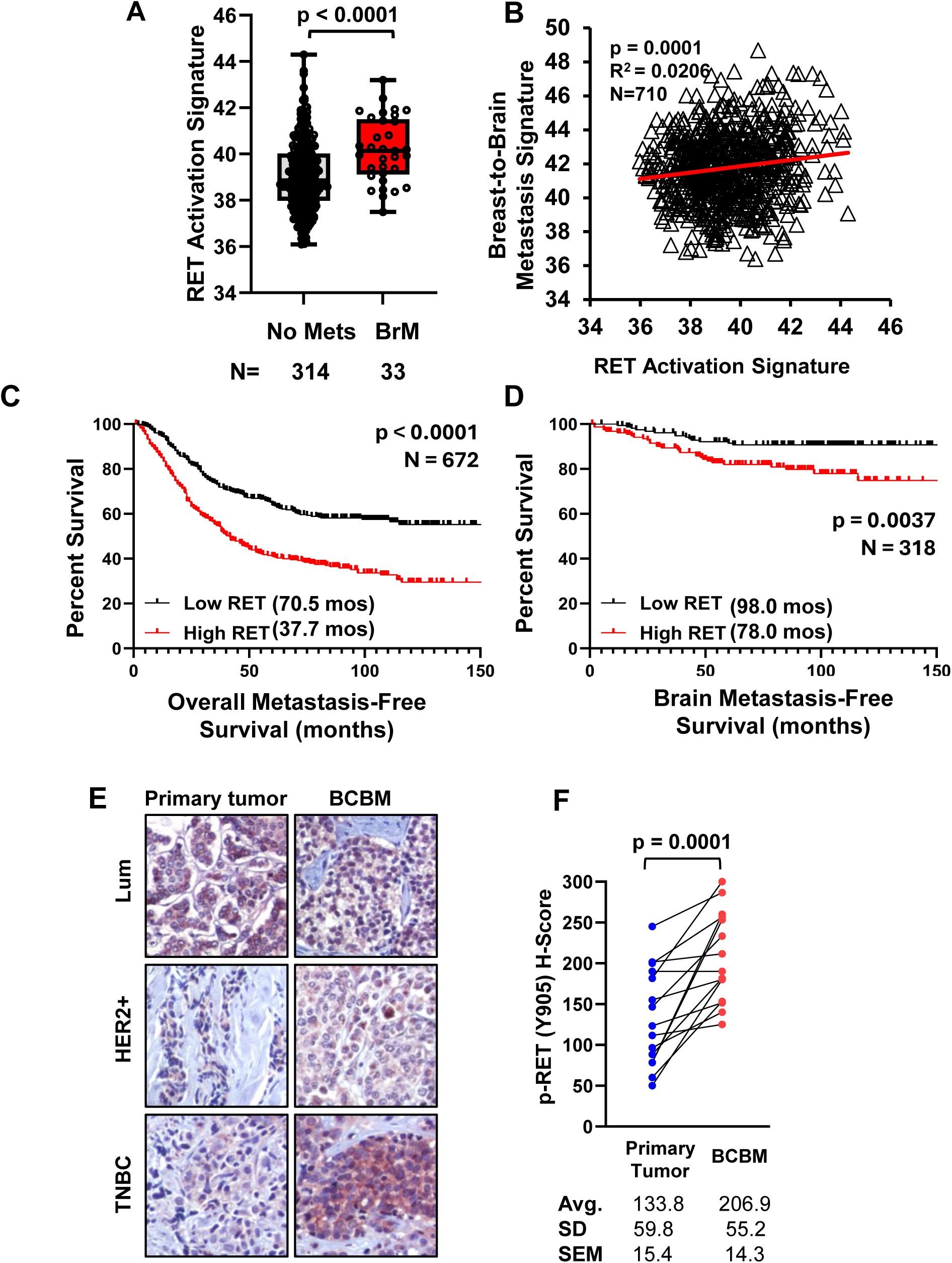
RET activity is enriched in BCBM and is associated with shorter overall and brain metastasis-free survival. (A) RET pathway activation scores between breast cancer patients with no metastases vs. brain metastases using GEO breast cancer datasets (GSE2034, GSE2603, GSE5327, GSE12276, GSE14020). An unpaired, two-tailed t-test was used in panel (A). (B) Pearson correlation analysis using GEO correlating RET activation signatures to a breast-to-brain metastasis signature. (C) Using the Kaplan-Meier analysis, log-rank analyses, GEO datasets, and the RET activation signatures, 672 breast cancer patients were stratified into two groups, either a high or low RET pathway activation for overall metastasis-free survival. Median of overall metastasis-free survival times are indicated in months (N=672). (D) High versus low RET activation scores were analyzed for brain metastasis-free survival (N = 318). (E) IHC staining of p-RET (Y905) expression levels in 15 matched pairs of primary breast tumors and their corresponding BCBM samples either luminal, HER2-enriched, or TNBC subtypes (20X images presented). (F) H-score quantification of p-RET (Y905) positivity, presented as a ladder plot (N=15). A paired, two-tailed t-test was used.

### RET activation is enriched in brain-topic HER2-enriched and TNBC cells and is associated with shorter brain metastasis-free survival in TNBC patients

Given that RET activation is enriched in BCBM patient samples, we next sought to determine whether this activation is recapitulated in brain-tropic breast cancer cell lines, enabling *in vitro* investigation of the pathway. We performed Western blot analyses using three pairs of parental breast cancer cells and their corresponding brain-tropic derivatives: HER2-enriched SKBR3 and brain-tropic SKBRM, as well as triple-negative CN34, CN34-BrM, and MDA-MB-231 and MDA-MB-231-BrM. These cell lines were selected not only for their availability as brain-tropic sublines but also due to their HER2-enriched or triple-negative subtypes, which are known to have a higher propensity for brain metastasis in breast cancer patients [2–4]. We found that across all three pairs of cell lines, the brain-tropic breast cancer cells demonstrated robustly higher expression of p-RET (Y905) when compared to their parental counterparts, further indicating that RET activity is preferentially elevated in brain-metastatic breast cancer cells (Fig. 2A). To assess downstream signaling of RET, we also evaluated the activation status of key effectors from the AKT and ERK pathways, both of which are downstream of RET [8]. Consistently, we found that the brain-tropic breast cancer cells also had increased expression levels of p-AKT (S473) and p-ERK (T202/Y204), further supporting enhanced RET signaling in these brain-tropic cells. These findings suggest that RET activation may play a particularly relevant role in BCBM arising from HER2-enriched and triple-negative breast cancer subtypes. To further investigate the clinical relevance of these observations, we conducted data mining analysis using our curated GEO breast cancer patient’s dataset and analyzed RET activation across breast cancer subtypes. Our analysis revealed that RET activation was significantly enriched in TNBC patients compared to HER2-enriched or luminal subtypes (Fig. S1A). Then, we stratified our survival analyses (Fig. 2C–D) by breast cancer subtype: triple-negative, HER2-enriched, or luminal breast cancer. We found that within all subtypes, patients with high RET activation had significantly shorter overall metastasis-free survival compared to those with low RET activation (Fig. 2B and D and F). Notably, among TNBC patients, we found that high RET activation was also significantly associated with shorter brain metastasis-free survival (Fig. 2C). In contrast, HER2-enriched breast cancer patients showed no significant differences in brain metastasis-free survival based on RET activation status (Fig. 2E), though this may be attributed to the small sample size of HER2-enriched patients with brain metastases in this dataset. Similarly, no significant differences in brain metastasis-free survival were observed among luminal breast cancer patients (Fig. 2G).

**Figure 2:**
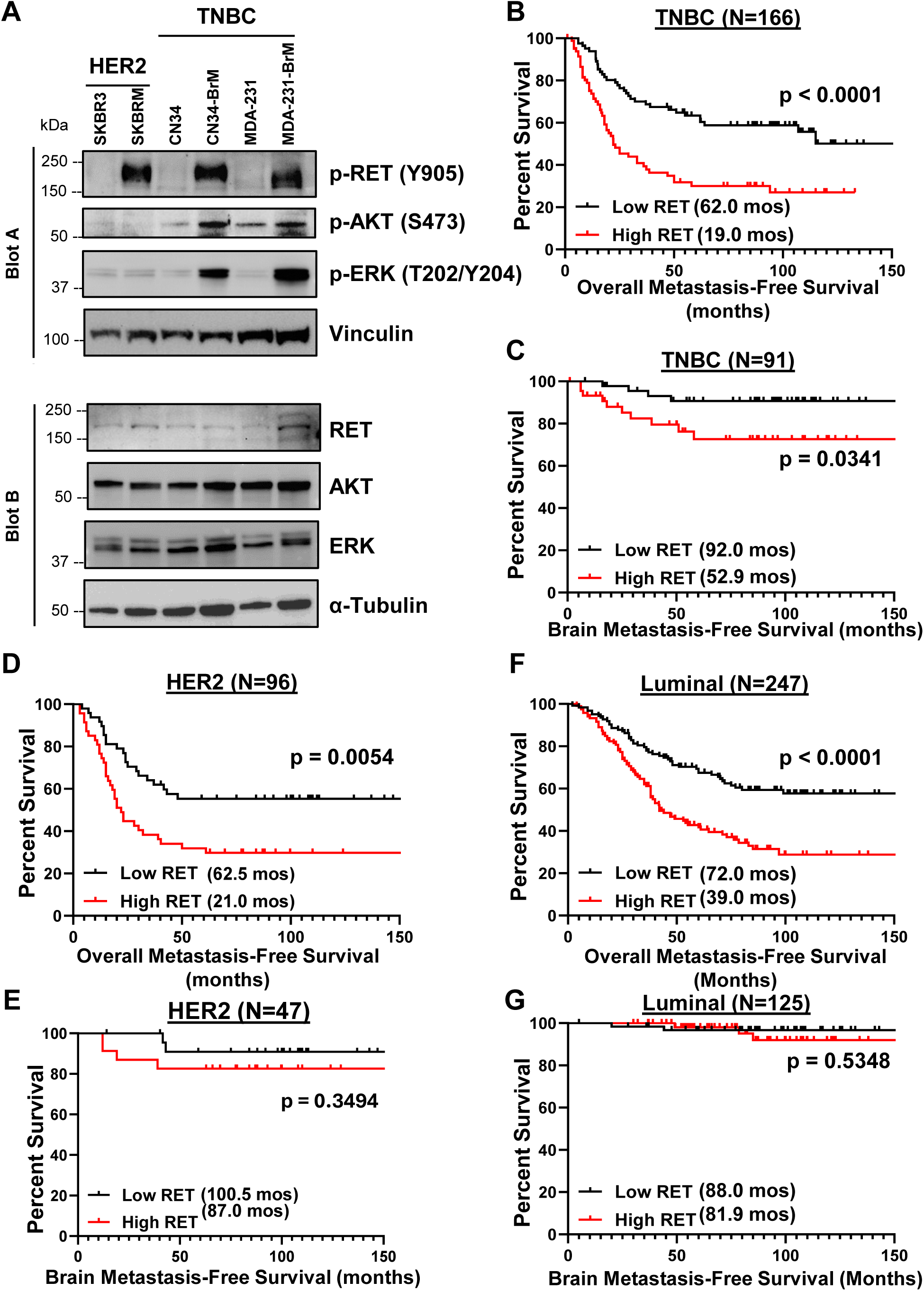
RET is frequently expressed and activated in brain-tropic breast cancer cells and is associated with shorter overall metastasis-free survival across subtypes and shorter brain metastasis-free survival in TNBC. (A) Western blot panel of three pairs of parental breast cancer cells and their corresponding brain-tropic derivatives to determine endogenous p-RET, p-AKT, p-ERK, total RET, total AKT, and total ERK protein expression. Vinculin serves as a high molecular weight loading control, and α-tubulin serves as a low molecular weight loading control. Kaplan-Meier analysis using high versus low RET activation scores were analyzed for patients with (B) TNBC metastasis-free survival (N = 166) and (C) brain metastasis-free survival (N = 91), (D) HER2-enriched metastasis-free survival (N = 96) and (E) brain metastasis-free survival (N = 47), and (F) luminal metastasis-free survival (N = 247) and (E) brain metastasis-free survival (N = 125) using the curated GEO breast cancer dataset.

### Overexpression of RET enhances brain-tropism of breast cancer cells in an intracardiac model of metastasis

Given the significant role our findings demonstrate for RET activation in BCBM, we sought to investigate whether RET overexpression can enhance brain tropism of parental breast cancer cells. We generated stable RET-overexpressing cells, MDA-MB-231-RET, from the parental MDA-MB-231 line. We performed intracardiac inoculation of female athymic nu/nu mice using MDA-MB-231-RET cells, followed by bioluminescent imaging twice weekly beginning seven days post-inoculation and continuing until the study endpoint at 28 days, at which mice were euthanized, and organs were collected for *ex vivo* analysis (Fig. 3A). RET overexpression was confirmed via Western blot, which showed robust expression of total RET and p-RET (Y905) (Fig. 3B). *In vivo* bioluminescent imaging revealed that RET-overexpressing breast cancer cells developed a significantly higher overall metastatic burden compared to the control breast cancer cells (Fig. 3C). *Ex vivo* analyses of resected organs at study endpoint demonstrated a significantly higher brain metastasis burden in mice inoculated with RET-overexpressing cells relative to control (Fig. 3D and E). Although trends towards increased metastases in bone (Fig. 3F) and liver (Fig. 3G) were observed, these differences were not statistically significant. Lastly, we assessed changes in body weights as a proxy for health status. We found that mice inoculated with RET-overexpressing cells exhibited significant weight loss over time, consistent with a higher metastatic burden and deteriorating health (Fig. 3H). Together, these findings demonstrate that RET overexpression promotes brain-tropism of TNBC cells *in vivo*.

**Figure 3:**
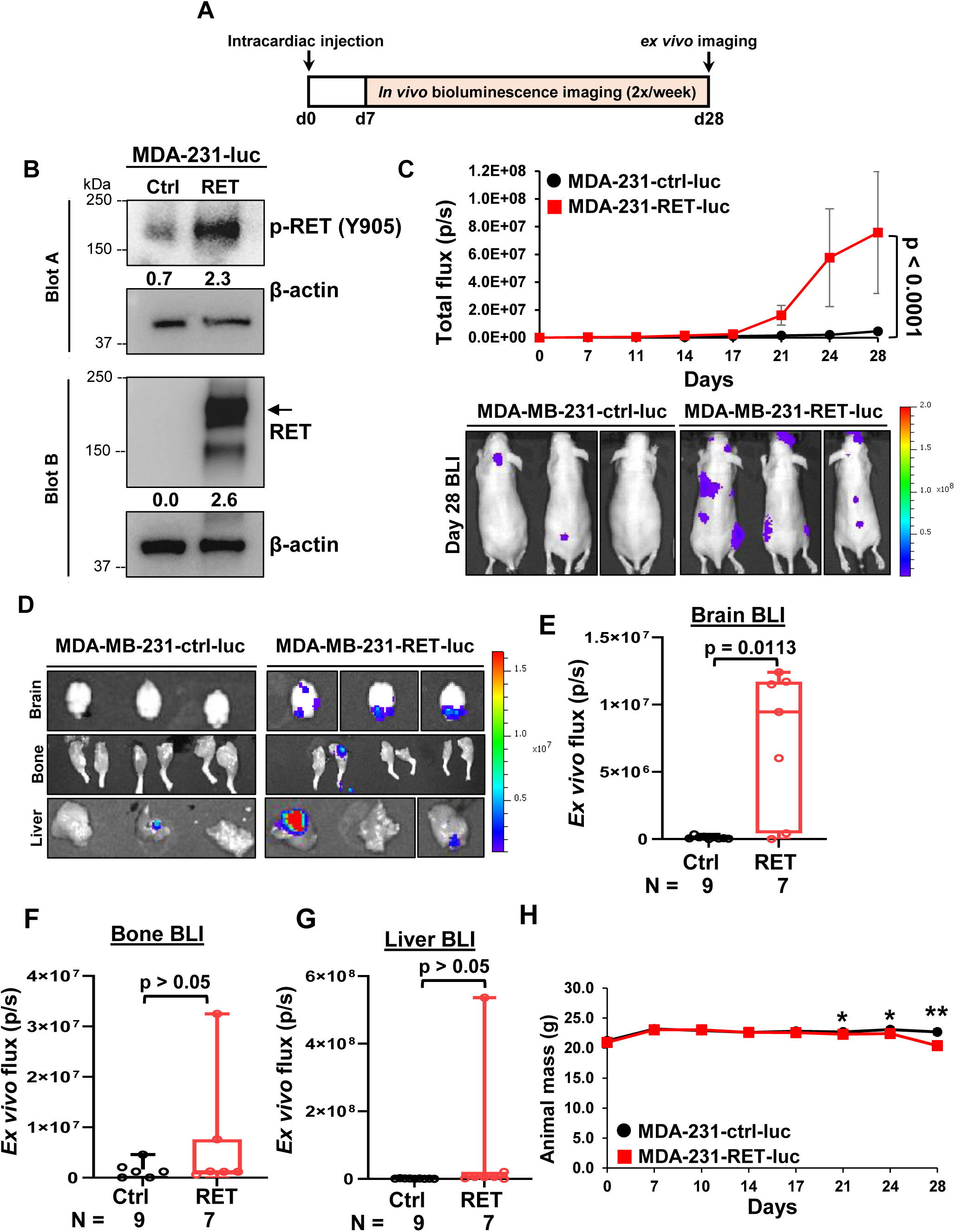
RET overexpression promotes brain metastasis of TNBC cells using an intracardiac mouse model. (A) Schematic of intracardiac mouse model. 6–7-week-old athymic nude mice were inoculated via the left ventricle with luciferase-expressing MDA-MB-231-RET cells (N=7) or MDA-MB-231-Ctrl cells(N=9). Mice were imaged twice weekly to monitor metastasis formation and growth. At the study endpoint (Day 28), mice were euthanized, and organs were collected for *ex vivo* analyses. (B) Confirmation of MDA-MB-231cell lines stably expressing control vector (Ctrl), or RET-expression vectors (RET) Western blot analysis. (C) Weekly average metastatic burden per group throughout the study, assessed by BLI. Representative BLI images from Day 28 (bottom). (D) Representative BLI images of *ex vivo* brain, bone, and liver metastases. Quantification of *ex vivo* BLI of (E) brains, (F) bones, and (G) livers, (H) Average weight of mice in each group. Two-way ANOVAs were used to calculate p-values with a mixed-effects analysis for panels C and H. An unpaired, two-tailed t-test was used in panels E, F and G.

### RET overexpression enhances intracranial growth of breast cancer cells

Building on our observation of a significant increase in metastatic burden following intracardiac injection of RET-overexpressing cells, we next sought to determine whether RET overexpression enhances the growth of established breast cancer brain metastases. To address this, we performed stereotaxic intracranial injections of RET-overexpressing breast cancer cells into female athymic nu/nu mice. Bi-weekly bioluminescent imaging was conducted seven days after inoculation and continued until the study endpoint at 28 days, at which mice were euthanized, and organs were collected for *ex vivo* analysis (Fig. 4A). We found that RET-overexpressing TNBC cells had significantly higher tumor burden in the brain when compared to control cells (Fig. 4B). *Ex vivo* analysis of mouse brains confirmed these findings, with RET-overexpressing tumors exhibiting significantly higher brain tumor burdens than those from control mice (Fig. 4C). *Ex vivo* IHC analysis of mouse brain tumors revealed markedly elevated levels of p-RET in RET overexpressing mice (Fig. 4D and E). Furthermore, mice inoculated with RET-overexpressing cells exhibited significantly higher tumoral proliferation, as evidenced by higher Ki-67 expression compared to control mice, indicating that RET overexpression promotes the tumor growth of established BCBM. While intracranial injections directly deliver tumor cells into the brain, tumor retention rates are often lower than anticipated. Consistent with this, our study found that 100% (12/12) of mice inoculated with RET-overexpressing breast cancer cells had detectable brain tumors, which is significantly higher than the 56% (6/11) observed in control mice, suggesting that RET overexpression substantially enhances breast cancer cell growth in the brain (Fig. 4F). No significant differences in average body weight were observed throughout the study (Fig. 4G).

**Figure 4:**
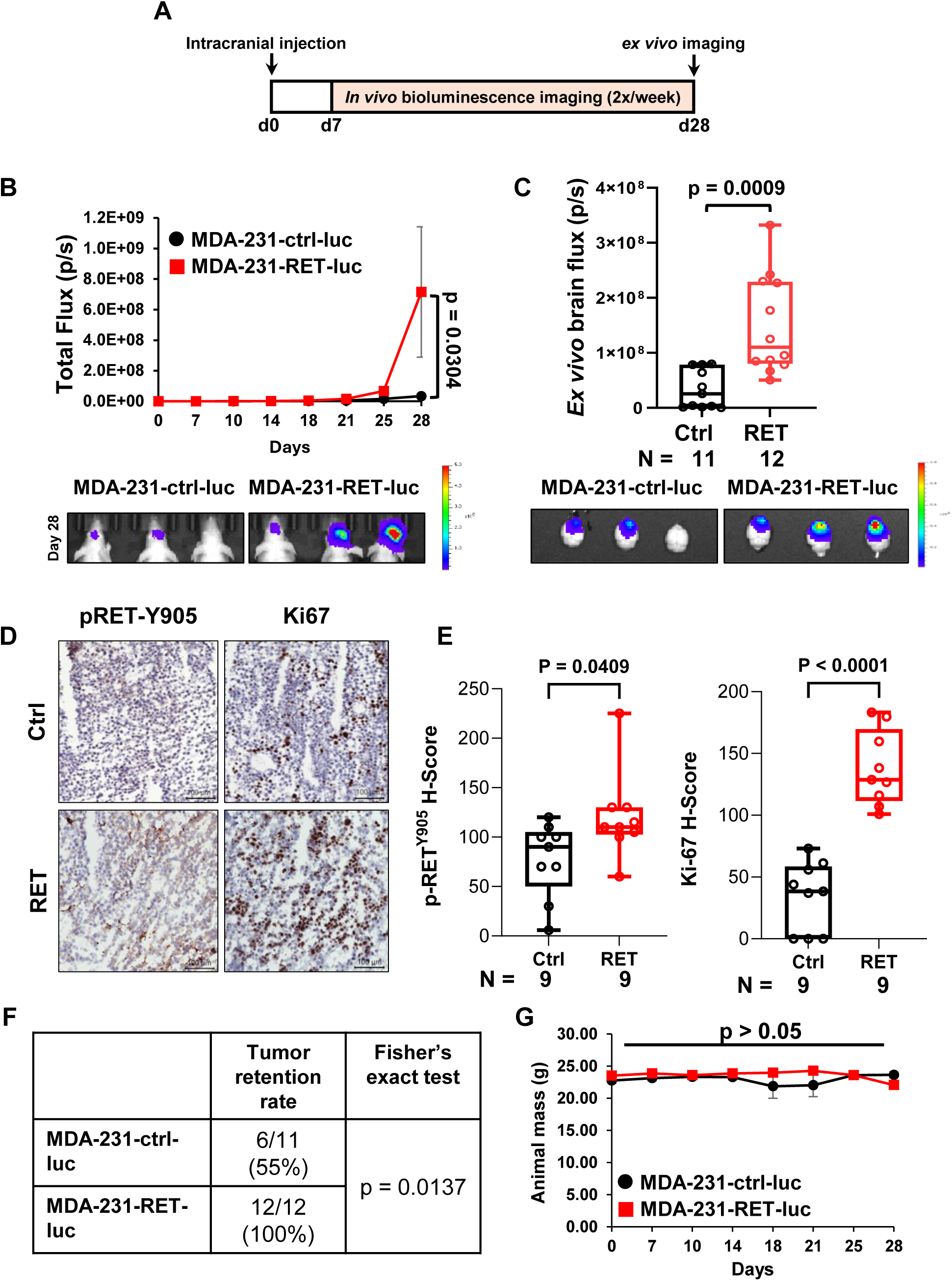
RET overexpression promotes tumor growth and proliferation of TNBC cells in the brain using an intracranial mouse model. (A) Schematic of intracranial mouse model. 6–7-week-old athymic nude mice were intracranially inoculated with luciferase-expressing MDA-MB-231-RET cells (N=12) or MDA-MB-231-Ctrl cells (N=11). Mice were imaged twice weekly to monitor tumor formation and growth. At the study endpoint (Day 28), mice were euthanized, and organs were collected for *ex vivo* analyses. (B) Bi-weekly average brain tumor burden per group throughout the study, assessed by BLI (top). Representative BLI images from Day 28 (bottom). (C) Quantification of *ex vivo* BLI of brains (top). Representative BLI images of *ex vivo* brain (bottom). (D) IHC staining of p-RET (Y905) and Ki-67 expression levels in brain tumors (scale bar 100µm). (E) H-score quantification of p-RET (Y905) positivity and Ki-67 positivity. (F) Tumor retention rate following inoculation. (H) Average weight of mice in each group. Two-way ANOVAs were used to calculate p-values with a mixed-effects analysis for panels (B and G). An unpaired, two-tailed t-test was used in panels (C and E).

### Selective inhibition of RET potently reduces cell viability and enhances apoptosis in brain-tropic breast cancer cell lines

To assess the therapeutic potential of selective RET inhibition in BCBM, we used three pairs of parental breast cancer cells and their brain-tropic variants. We treated these cell lines with increasing doses of either pralsetinib or selpercatinib, and the half-maximal inhibitory concentration (IC50) was determined for each treatment. Our results show that breast cancer cells, both parental and brain-tropic variants, were sensitive to selective RET inhibition, with no significant differences in sensitivity between the two groups (Fig. S2A). Treatment with 25µM of pralsetinib significantly reduced cell viability of SKBRM (35%), CN34-BrM (51%), and MDA-MB-231-BrM (42%) compared to vehicle-treated cells (Fig. 5A). Similarly, 25µM selpercatinib also significantly reduced cell viability of all three brain-tropic cell lines (Fig. 5B), suggesting that selective RET inhibition may be a viable therapeutic strategy for targeting brain-tropic breast cancer cells. To examine changes in signaling activity following RET inhibition, we performed Western blot analysis on brain-tropic breast cancer cells treated with pralsetinib or selpercatinib. We found that treatment with 10 µM pralsetinib (Fig. 5C, left panel) potently inhibited RET activity (p-RET Y905) and downstream signaling, including STAT3 (p-STAT3 Y705), and ERK (p-ERK 1/2). While treatment with selpercatinib also significantly inhibited RET activation, we observed only a modest reduction in STAT3 activation and ERK phosphorylation (Fig. 5C, right panel). Given the significant reduction in cell viability, we next sought to determine whether selective RET inhibition induces apoptosis in brain-tropic breast cancer cells. SKBRM, CN34-BrM, and MDA-231-BrM were treated with vehicle, pralsetinib (10µM), or selpercatinib (10µM), and Annexin V flow cytometry was performed. Annexin V binds to calcium-dependent phospholipids exposed to the extracellular space during cellular stress, and increased Annexin V staining correlates with increased apoptosis [29]. We observed significantly higher Annexin V positivity in CN34-BrM (Fig. 5E) and MDA-231-BrM (Fig. 5F) brain-tropic cell lines following treatment with pralsetinib, and significantly higher Annexin V positivity in all brain-tropic breast cancer cell lines used (Fig. 5D and E) following treatment with selpercatinib, indicating enhanced apoptosis upon selective inhibition of RET. Additionally, we assessed cleaved poly-(ADP ribose) polymerase (PARP) as an alternative marker for apoptosis [30]. Full-length PARP is a DNA damage sensor that is cleaved into fragments (cleaved PARP) during caspase-dependent apoptosis. Both pralsetinib (Fig. 5G) and selpercatinib (Fig. 5H) treatments induced cleaved PARP expression compared to vehicle controls, with selpercatinib treatment resulting in a more pronounced increase in cleaved PARP levels.

**Figure 5:**
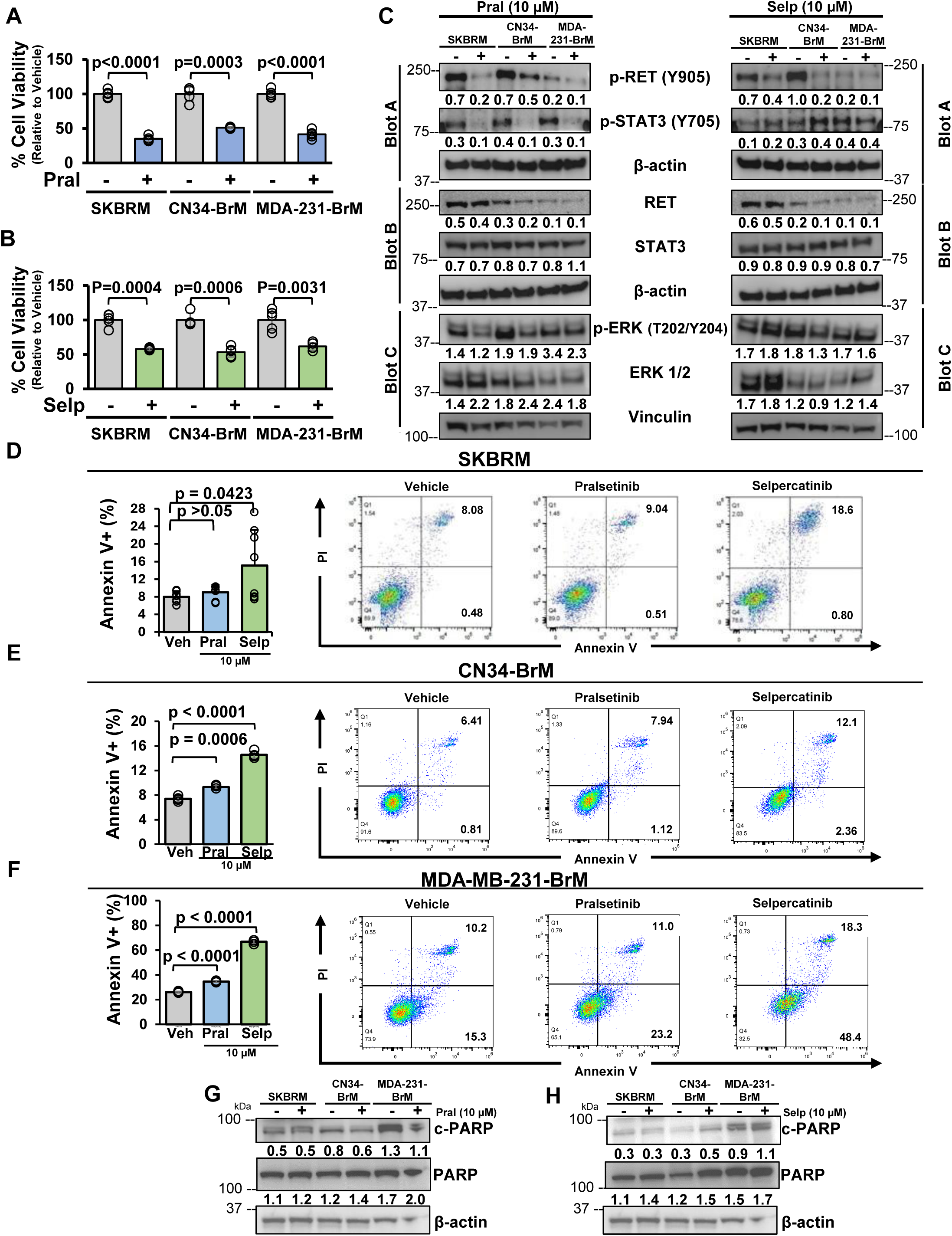
RET inhibitors reduce cell viability, suppress downstream RET signaling, and induce apoptosis in brain-tropic breast cancer cells *in vitro*. The effect of treatment with (A) pralsetinib or (B) selpercatinib on cell viability of brain-tropic breast cancer cell lines (N=5). (C) Western blot analysis to determine protein expression of p-RET (Y905), p-STAT3 (Y705), total RET, total STAT3, p-ERK (T202/Y204), and total ERK ½ following treatment with prasetinib (left) or selpercatinib (right). Vinculin serves as high molecular weight loading control, and β-actin serves as a low molecular weight loading control. Annexin V/PI flow cytometry co-staining of (D) SKBRM cells, (E) CN34-BrM cells, and (F) MDA-MB-231-BrM cells treated with pralsetinib or selpercatinib (quantification on the left and representative images on the right) (N=5). Western blot analysis of cleaved PARP and full-length PARP in brain-tropic breast cancer cells following treatment with (G) pralsetinib, or (H) selpercatinib. An unpaired, two-tailed t-test between vehicle and treatment was used in panels (A,B,D,E and F).

### RET inhibition impairs the migratory capacity of brain-tropic breast cancer cells *in vitro*

Migratory capacity is a critical step in the metastatic cascade. We therefore investigated whether pharmacological inhibition of RET could impair the migration of brain-tropic breast cancer cells *in vitro*. We performed scratch wound assays using SKBRM, CN34-BrM, and MDA-MB-231-BrM cell lines treated with vehicle, pralsetinib, or selpercatinib. Images were acquired immediately following scratching and treatment administration (T0), and again 24 hours later (T24). We found that in SKBRM cells, treatment with either pralsetinib (Fig. 6A) or selpercatinib (Fig. 6B) significantly reduced cell migration compared to vehicle treatment. Similar results were observed in CN34-BrM (Fig. 6C and D) and MDA-MB-231-BrM cells (Fig. 6E and F), indicating that RET inhibition effectively impairs migratory ability across multiple brain-tropic breast cancer cell lines. To determine whether RET inhibition also impairs the invasive capacity, a key step for metastatic colonization, of brain-tropic breast cancer cell lines, we performed basements membrane invasion assays in the presence of pralsetinib or selpercatinib for 24h. In contrast to our migration data, we found that the invasion of SKBRM (Fig. 6G) and CN34-BrM (Fig. 6H) cells was not significantly affected by treatment with RETi. While MDA-MB-231-BrM cells demonstrated a trend toward reduced invasion following treatment with either pralsetinib or selpercatinib (Fig. 6I), these changes did not reach statistical significance. The lack of significant effects on invasion suggests that RET plays a more dominant role in regulating migration rather than invasion in brain-tropic breast cancer cells *in vitro*. Together, our data suggests that selective pharmacological inhibition of RET effectively impairs cell migration, but not invasion, in brain-tropic breast cancer cells *in vitro*.

**Figure 6:**
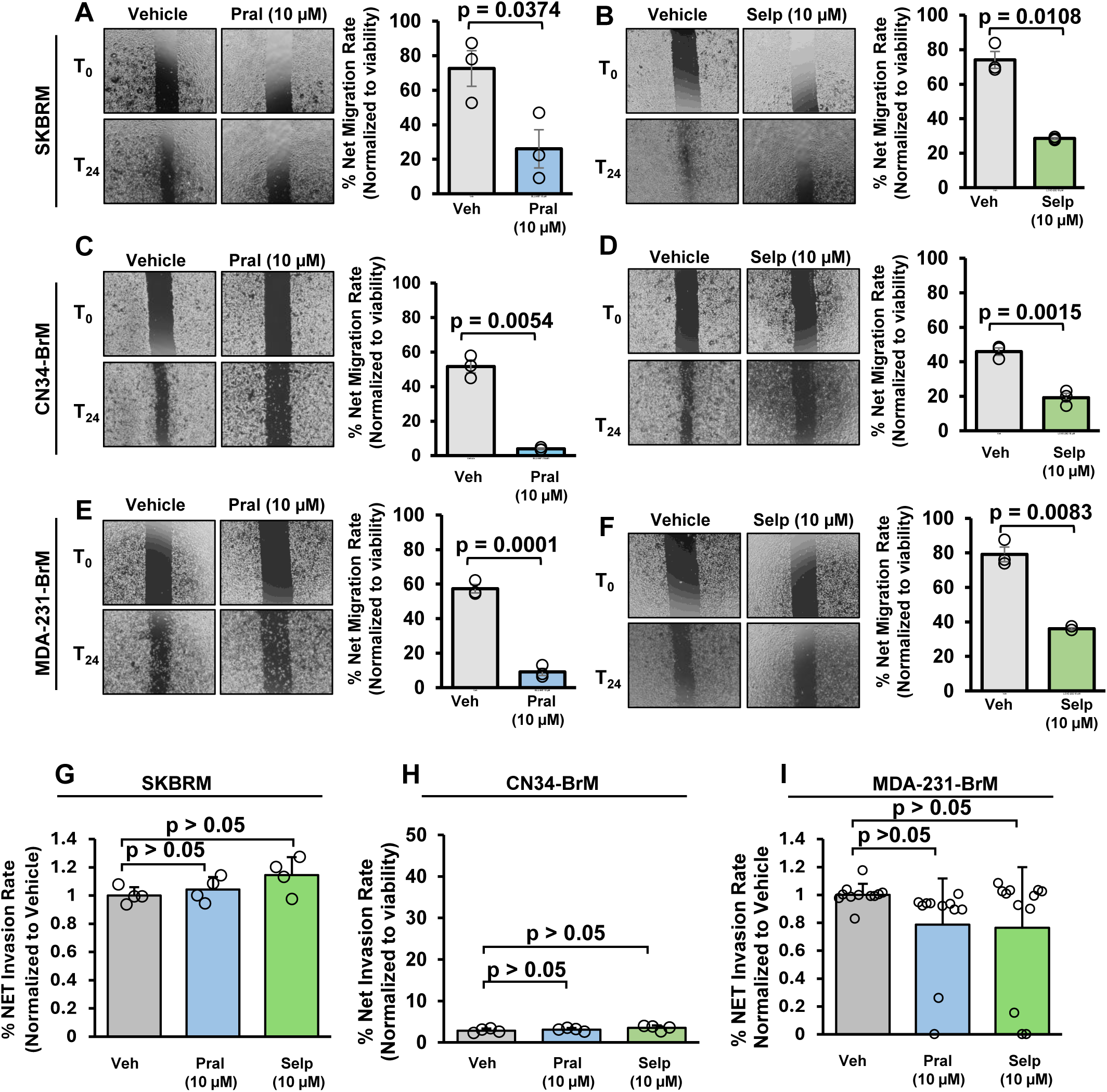
RET inhibition impairs the migratory capacity of brain-tropic breast cancer cells *in vitro*. Net migration rate of brain-tropic breast cancer cells after 24-h treatment of vehicle of pralsetinib (A,C, and E) of selpercatinib (B,D, and F) (N=3). (G)Net migration rate of SKBRM cells (N=4), (H) CN34-BrM cells (N=4), and (I) MDA-MB-231-BrM cells (N=12) following treatment with pralsetinib or selpercatinib. An unpaired, two-tailed t-test between vehicle and treatment was used in all panels.

### Pharmacological inhibition of RET, using selpercatinib, reduces brain metastasis in a preventative model of systemic breast cancer metastasis *in vivo*

Our findings thus far demonstrate that RET inhibition potently inhibits brain-tropic breast cancer cell viability, induces apoptosis, and impairs migratory capacity *in vitro*, warranting the evaluation of the therapeutic efficacy of RET inhibition against breast cancer metastasis *in vivo*. To this end, we utilized the brain-tropic TNBC cells MDA-MB-231-BrM in two intracardiac inoculation-based metastasis models in female athymic nu/nu mice: (1) a preventative (pre-brain) metastasis model, in which daily oral administration of selpercatinib began one day after tumor cell inoculation, and (2) a treatment (post-brain) metastasis model, in which daily selpercatinib administration was initiated only after brain metastases where confirmed via bioluminescent imaging (Fig. 7A). In our preventative (pre-brain) metastasis model, we found that daily oral administration of selpercatinib significantly reduced brain metastasis burden compared to vehicle treated mice (Fig. 7B). In contrast, selpercatinib treatment did not confer significant therapeutic benefit in the treatment (post-brain) metastasis model (Fig. 7C), suggesting that selective RET inhibition may yield a more effective therapeutic benefit when administered earlier in the metastatic process, prior to overt colonization of the brain. *Ex vivo* analyses of resected organs (Fig. 7D) supported these findings. Mice from our preventative (pre-brain) metastasis model treated with selpercatinib had significantly lower brain (Fig 7E) and bone (Fig. 7F) metastasis when compared to vehicle treated mice. However, our treatment (post-brain) model showed no significant differences in metastatic burden of the tissues between treatment groups. Liver metastases were unaffected by selpercatinib treatment in either model (Fig. 7G). Given that selpercatinib is predominantly metabolized in the liver by CYP3A4, we assessed potential hepatotoxicity by measuring serum alanine transaminase (ALT) activity *ex vivo* [31]. We found no significant differences in ALT levels between groups, and values remained well below the reported toxicity threshold (109 ± 18 U/L) (Fig. 7H). Additionally, we monitored animal weights throughout the study as an indicator of general health and systemic toxicity. There were no significant changes in average body weight were detected in any treatment group (Fig. 7I and J), further supporting the tolerability of selpercatinib in both models. Together, these findings provide strong evidence that early intervention with RETi can significantly impair brain metastasis formation.

**Figure 7:**
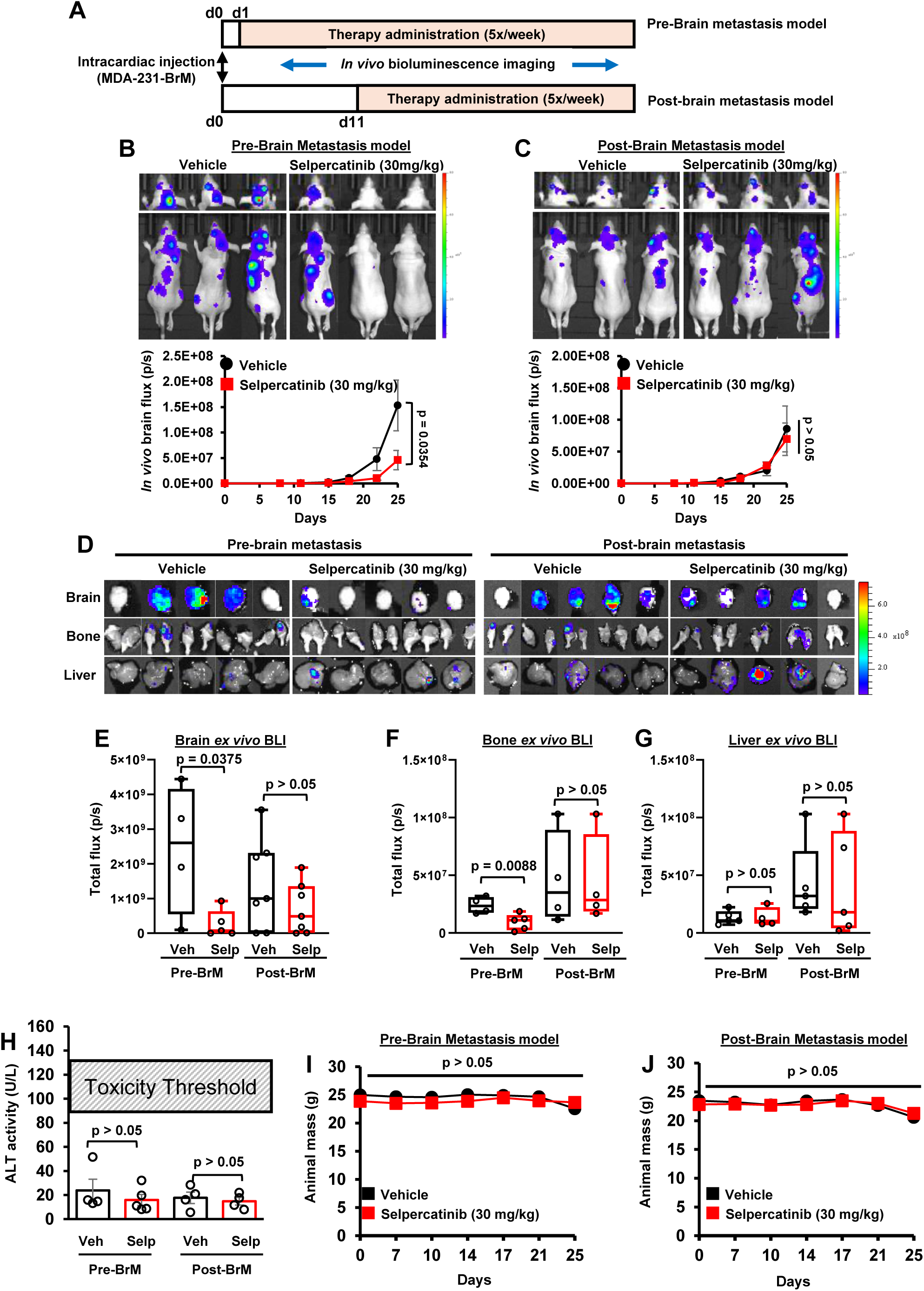
Systemic administration of selpercatinib selectively reduces brain metastasis formation *in vivo*. **(A) Schematic of intracardiac mouse model.** 6–7-week-old athymic nude mice were inoculated via the left ventricle with luciferase-expressing MDA-MB-231-BrM cells. Mice (N=5/group) in pre-brain metastasis model (top) received first treatment 1 day after inoculation, and mice (N=8/group) in the post-brain metastasis model received treatment 11 days after inoculation. Mice were treated 5 times a week and imaged twice weekly to monitor metastasis formation and growth. At the study endpoint (Day 25), mice were euthanized, and organs were collected for *ex vivo* analyses. Weekly average metastatic burden per group throughout the study, assessed by BLI (bottom), and representative BLI images from Day 25 (top) of (B) pre-brain metastasis model and (C) post-brain metastasis model. (D) Representative BLI images of *ex vivo* brain, bone, and liver metastases. Quantification of *ex vivo* BLI of (E) brains, (F) bones, and (G) livers. (H) Circulating ALT levels between treatment groups. The average weight of mice in (I) pre-brain metastasis and (J) post-brain metastasis groups. Two-way ANOVAs were used to calculate p-values with a mixed-effects analysis for panels (B, C, I and J). An unpaired, two-tailed t-test was used in panels (E, F, G and H).

### Pralsetinib treatment reduces brain metastatic tumor burden and enhances tumor cell apoptosis *in vivo*

Given the therapeutic efficacy of selpercatinib in our preventative metastasis model, we next sought to investigate whether pralsetinib inhibits brain colonization in brain metastasis model *in vivo*. Here we employed an intracranial inoculation-based model, in which daily oral pralsetinib treatment was initiated two days after intracranial injection of the brain-tropic TNBC cells MDA-MB-231-BrM (Fig. 8A). Consistent with our previous findings, we found that pralsetinib treatment significantly reduced brain tumor burden as assessed by *in vivo* bioluminescent imaging (Fig. 8B). These findings were corroborated by *ex vivo* quantification of brain tumors further confirming this reduction (Fig. 8C). Of note, pralsetinib is also metabolized in the liver via the cytochrome P450 system so we sought to evaluate potential liver toxicity [32]. We measured serum ALT levels and found no significant differences between treatment and control groups (Fig. 8D). Likewise, we also monitored animal weights throughout the study and observed no significant differences in body weight during the study period (Fig. 8E), suggesting overall tolerability of pralsetinib. To gain mechanistic insight into the effects of RET inhibition *in vivo*, we performed IHC analysis on brain tumor sections and found that pralsetinib-treated mice exhibited a marked reduction in p-RET expression, confirming effective RET pathway inhibition *in vivo* (Fig. 8F). Furthermore, there was a significant decrease in Ki-67 staining, indicating reduced tumor cell proliferation in response to treatment (Fig. 8F). To further assess the pro-apoptotic effects of RET inhibition, we conducted a TUNEL assay on brain sections. We observed a significant increase in TUNEL-positive cells with pralsetinib treatment compared to vehicle controls, suggesting that pralsetinib promotes tumor cell apoptosis in the brain metastatic niche *in vivo* (Fig. 8G). Collectively, these findings provide strong evidence that selective RET inhibition can significantly impair BCBM growth, reduce tumor proliferation, and enhance apoptosis, all while being well tolerated *in vivo*.

**Figure 8:**
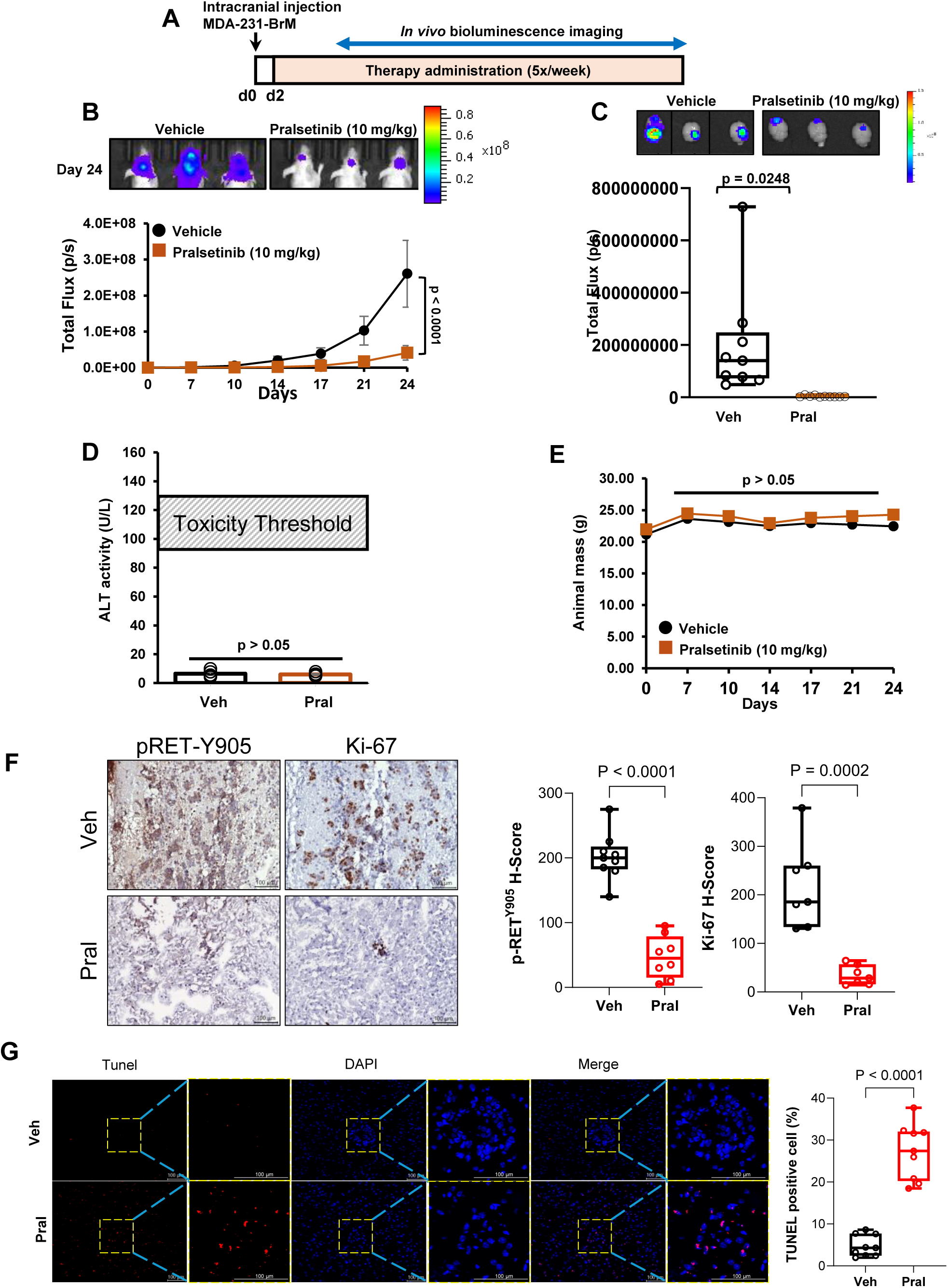
Systemic administration of pralsetinib selectively reduces established brain metastatic tumor growth and proliferation *in vivo*. (A) Schematic of intracranial inoculation mouse model. 6–7-week-old athymic nude mice (N=10/group) were inoculated with luciferase-expressing MDA-MB-231-BrM cells. Mice received their first treatment 2 days after inoculation and were treated 5 times a week and imaged twice a week to monitor tumor growth. At the study endpoint (Day 24), mice were euthanized, and brains were collected for *ex vivo* analyses. (B) Weekly average metastatic burden per group throughout the study, assessed by BLI (bottom), and representative BLI images from Day 25(top), vehicle N = 9 and Pralsetinib N = 10. (C) Representative BLI images of *ex vivo* brain (top). Quantification of *ex vivo* BLI of brains (bottom). (D) Circulating ALT levels between treatment groups. (E) The average weight of mice throughout the study. (F) IHC staining of p-RET (Y905) and Ki-67 expression levels in brain tumors (left) (scale bar = 100µm), and H-score quantification of p-RET (Y905) positivity and Ki-67 positivity (right). (G) Fluorescent TUNEL assay on mouse brain tumors. Images of representative fields show nuclei from control and treated tumors (left) (scale bar = 100µm). Two-way ANOVAs were used to calculate p-values with a mixed-effects analysis for panels (B and E). An unpaired, two-tailed t-test was used in panels (C, D, F and G).

## DISCUSSION

BCBM is associated with poor prognosis and a notably short median survival time [4, 5]. To date, BCBM treatment remains an immense therapeutic challenge [6, 7]. In addressing the challenges posed by BCBM, we have investigated for the first time that targeting RET kinase with FDA-approved selective RETis offers a promising therapeutic strategy for treating brain-metastatic breast cancers, including HER2-enriched and TNBC. In this study, we have made the following novel observations: RET activation is significantly higher in breast cancer patients with brain metastases compared to patients without, and RET activation is significantly correlated with shorter brain metastasis-free survival. Expression of activated RET is preferentially elevated in BCBM and brain-tropic breast cancer cells compared to the matched tumor or parental cells, respectively. Non-brain-metastatic breast cancer cells with stable RET overexpression demonstrate enhanced brain metastasis and intracranial tumor growth following intracardiac or intracranial inoculation, respectively. Brain-tropic breast cancer cells are sensitive to RETis and demonstrate significantly reduced cell viability, reduced cell migration, and increased apoptosis following treatment with either pralsetinib or selpercatinib. Lastly, daily oral administration of RETis significantly reduces brain metastasis and intracranial tumor growth following intracardiac or intracranial injection of brain-tropic breast cancer cells, respectively.

Selective RETis, selpercatinib and pralsetinib, have demonstrated strong efficacy in RET-altered NSCLC and thyroid cancers, including significant intracranial responses [9, 13, 14, 16]. While the use of RETis has been recently explored in luminal breast cancers, their role in HER2-enriched and triple-negative subtypes is not yet evaluated [33, 34]. Building on their efficacy in brain metastases of other RET-driven tumors, further evaluation of their therapeutic potential in BCBM is warranted.

Previous reports indicate that RET is overexpressed in brain metastases of patients with metastatic breast cancers [34]. However, it was only recently that RET activity was evaluated in brain metastases of luminal breast cancer patients [33]. Due to the lack of effective treatments in patients with brain-metastatic HER2-enriched and triple-negative breast cancers, we sought to evaluate whether RET would be a viable treatment target in these breast cancer subtypes as they are also at higher risk for developing brain metastases [2, 35]. In agreement with previous reports, we investigated that high RET activity is associated with BCBM signature genes, poor prognosis and low survival rate in BCBM patients. Notably, RET activation and its down-stream signals AKT-ERK significantly enriched in brain metastases of TNBC. We also found that overexpression of RET results in an increase in metastatic tumor burden upon intracardiac injection of cells, as well as increased tumor growth in the brain upon intracranial injection. Moreover, we found that treatment with either RET inhibitor significantly reduced breast cancer cell proliferation and enhanced apoptosis when compared to vehicle treatment. Importantly, daily oral administration of selpercatinib significantly reduced brain metastasis burden of brain-tropic MDA-MB-231-BrM cells following intracardiac injection. Daily oral administration with pralsetinib also significantly reduced tumor growth in an intracranial injection model of MDA-MB-231-BrM cells. Given that RET overexpression enhanced the metastatic phenotype of breast cancer cells, we posited that pharmacological inhibition of RET would inhibit migratory and invasive capacity of brain-tropic breast cancer cells. Our current study demonstrates that selective RET inhibition, with either pralsetinib or selpercatinib, significantly reduced the migratory capacity of brain-tropic TNBC cells *in vitro*. These findings may be one of the multiple mechanisms by which increased RET activity can drive brain metastasis of breast cancers *in vivo*.

As selective RET inhibition also enhanced apoptosis, as demonstrated by our Annexin V/PI flow cytometry and cleaved PARP Western blot analyses, future mechanistic studies are warranted to determine the pathways through which RET elicits increased metastasis of breast cancers. In our metastasis treatment models, we reported that selpercatinib significantly reduced brain metastasis of MDA-MB-231-BrM cells. Similarly, our intracranial treatment model demonstrated a robust reduction in brain tumor growth of MDA-MB-231-BrM cells following treatment with pralsetinib. The potent reduction in brain metastasis and intracranial tumor growth by RET monotherapy supports previous reports that both selpercatinib and pralsetinib are capable of penetrating the BBB [13, 16]. Interestingly, we found that treatment with selpercatinib also reduced bone metastasis in the intracardiac injection model, but only if administered one day after inoculation. This suggests that RET monotherapy also has the potential for use in treatment of bone metastases, though this would warrant further investigation as to its clinical utility. Importantly, though our *in vivo* models demonstrated potent efficacy of RET monotherapies in reducing brain metastasis and intracranial tumor growth, the effect of selective RET inhibition on overall survival remains unclear. Though our intracardiac and intracranial treatment models demonstrated significant therapeutic efficacy, our studies were designed such that the samples were collected on the day of study endpoint, which would not allow for appropriate evaluation of survival. A future treatment study to evaluate survival, where the primary endpoint is animal death, would be a more appropriate model for this type of analysis. A similar survival study could be performed to evaluate the effect of RET overexpression on metastasis-induced mortality.

In conclusion, we provide novel evidence that supports an important role that RET plays in promoting BCBM and demonstrates RET monotherapy, using FDA-approved selective RET inhibitors pralsetinib or selpercatinib, is an effective treatment strategy against brain metastases of breast cancers, including HER2-enriched and triple-negative breast cancers. We demonstrate that selective RET inhibition elicited significant anti-tumor efficacy with no hepatotoxicity. The data provides the preclinical rationale to clinically evaluate the efficacy of FDA-approved RET inhibitors in BCBM of HER2-enriched and triple-negative breast cancers.

## AUTHOR CONTRIBUTIONS

**Angelina Regua**: conceptualization, data curation, formal analysis, resources, methodology, writing – original draft, writing – review & editing, and visualization. **Shivani Bindal**: data curation, formal analysis, methodology, writing – original draft, writing – review & editing, and visualization. **Mariana Najar:** data curation, formal analysis, methodology, writing – original draft, writing – review & editing. **Phi-Long Tran:** data curation, formal analysis, writing – review & editing, and visualization. **Joshua Cha**: data curation, formal analysis, writing – review & editing, and visualization. **Syed Shams:** data curation, formal analysis. **Hui-Wen Lo**: conceptualization, funding acquisition, investigation, methodology, project administration, resources, supervision, validation, visualization, writing – review & editing. All authors have read and agreed to the published version of the manuscript.

## ACKNOWLEDGMENTS

The authors would like to acknowledge Drs. Fei Xing and Kounosuke Watabe for the SKBRM cells, Dr. Joan Massagué for the CN34-BRM cells, and Dr. Wensheng Li for the tumor samples. We acknowledge funding support for this study from NIH grants R01CA228137 (H-WL) and R21CA286225 (H-WL), DoD grants W81XWH-19-1-0072 (H-WL), W81XWH-20-1-0044 (H-WL), W81XWH-19-1-0753 (H-WL), and HT9425-24-1-0889 (H-WL), as well as, MetaVivor Translational Research Grants (H-WL) and UTStars (H-WL).

Supplementary Figures and Original Western Blots are included.

## Original WB Images

**Fig. 2A.**
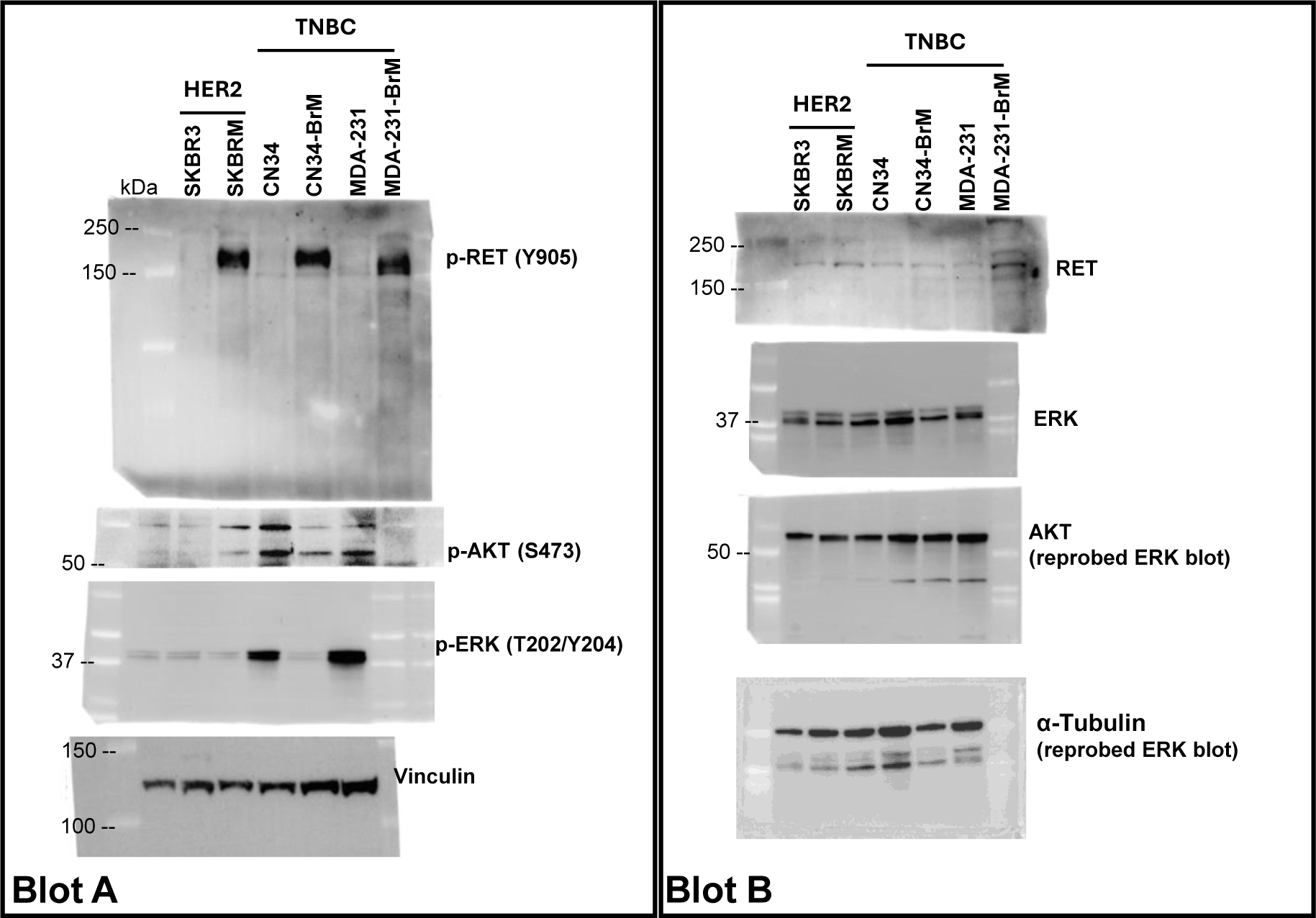

**Fig. 5C.**
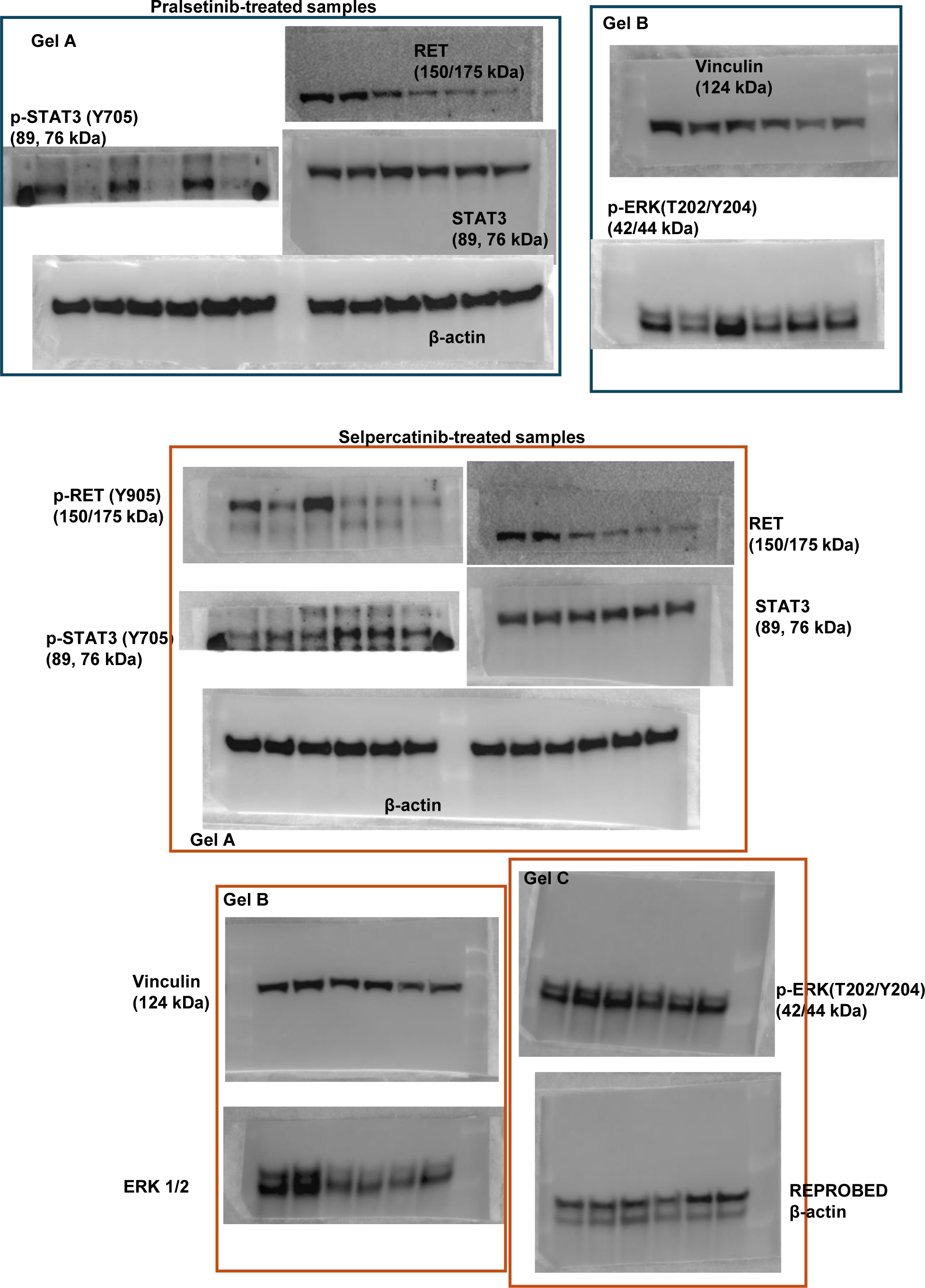

**Fig. 5G/5H.**
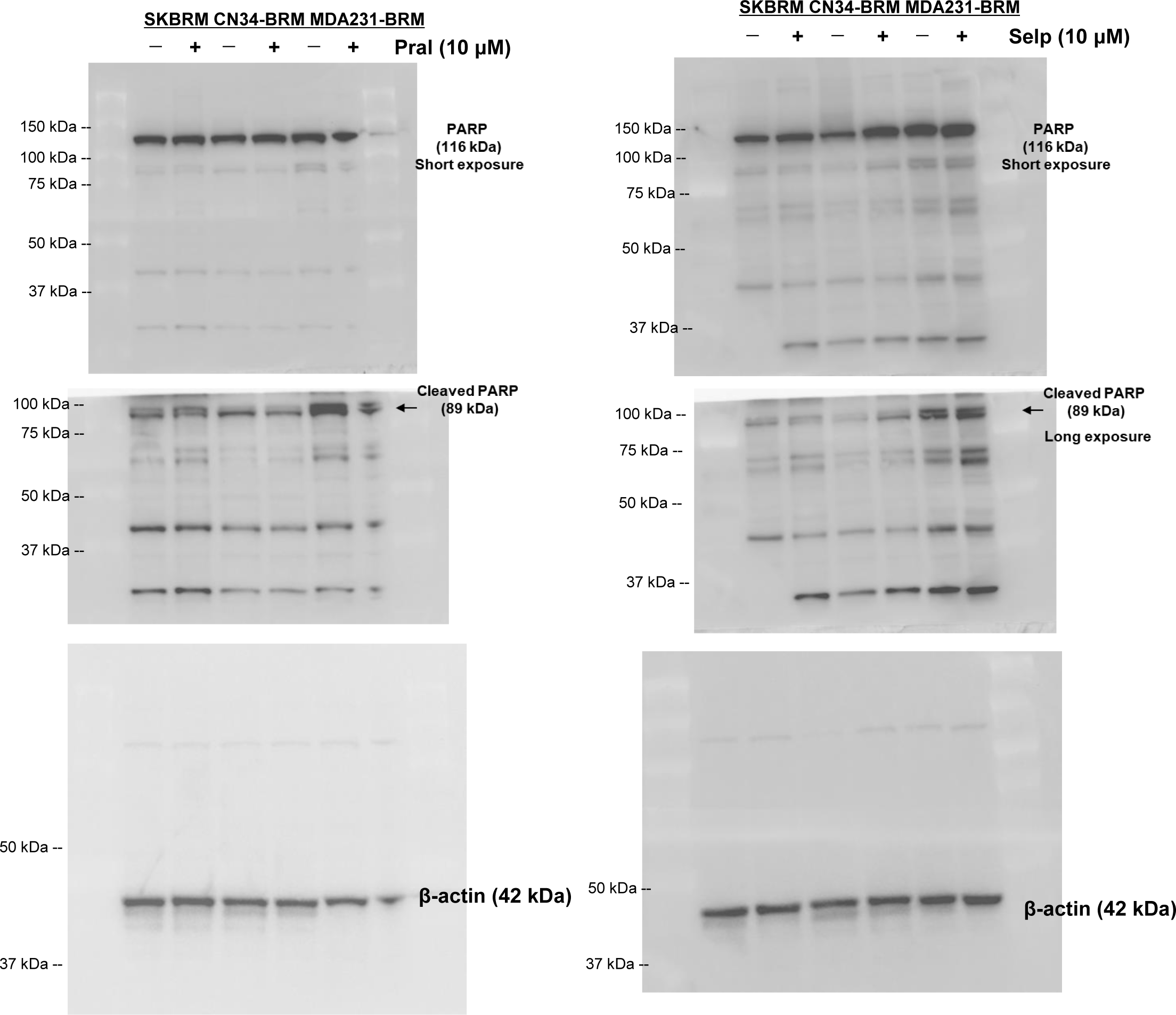

## REFERENCES

[1] R.L. Siegel, A.N. Giaquinto, A. Jemal, Cancer statistics, 2024, CA Cancer J Clin, 74 (2024) 12–49.

[2] A. Soni, Z. Ren, O. Hameed, D. Chanda, C.J. Morgan, G.P. Siegal, S. Wei, Breast cancer subtypes predispose the site of distant metastases, Am J Clin Pathol, 143 (2015) 471–478.

[3] Y.J. Kim, J.S. Kim, I.A. Kim, Molecular subtype predicts incidence and prognosis of brain metastasis from breast cancer in SEER database, J Cancer Res Clin Oncol, 144 (2018) 1803–1816.

[4] A. Darlix, G. Louvel, J. Fraisse, W. Jacot, E. Brain, M. Debled, M.A. Mouret-Reynier, A. Goncalves, F. Dalenc, S. Delaloge, M. Campone, P. Augereau, J.M. Ferrero, C. Levy, J.D. Fumet, I. Lecouillard, P. Cottu, T. Petit, L. Uwer, C. Jouannaud, M. Leheurteur, V. Dieras, M. Robain, M. Chevrot, D. Pasquier, T. Bachelot, Impact of breast cancer molecular subtypes on the incidence, kinetics and prognosis of central nervous system metastases in a large multicentre real-life cohort, Br J Cancer, 121 (2019) 991– 1000.

[5] I. Witzel, E. Laakmann, R. Weide, T. Neunhoffer, T.J. Park-Simon, M. Schmidt, P.A. Fasching, T. Hesse, A. Polasik, S. Mohrmann, F. Wurschmidt, C. Schem, C. Bechtner, R. Wurstlein, T. Fehm, V. Mobus, N. Burchardi, S. Loibl, V. Muller, Treatment and outcomes of patients in the Brain Metastases in Breast Cancer Network Registry, Eur J Cancer, 102 (2018) 1–9.

[6] N. Niikura, N. Hayashi, N. Masuda, S. Takashima, R. Nakamura, K. Watanabe, C. Kanbayashi, M. Ishida, Y. Hozumi, M. Tsuneizumi, N. Kondo, Y. Naito, Y. Honda, A. Matsui, T. Fujisawa, R. Oshitanai, H. Yasojima, Y. Tokuda, S. Saji, H. Iwata, Treatment outcomes and prognostic factors for patients with brain metastases from breast cancer of each subtype: a multicenter retrospective analysis, Breast Cancer Res Treat, 147 (2014) 103–112.

[7] R. Rostami, S. Mittal, P. Rostami, F. Tavassoli, B. Jabbari, Brain metastasis in breast cancer: a comprehensive literature review, J Neurooncol, 127 (2016) 407–414.

[8] A.T. Regua, M. Najjar, H.W. Lo, RET signaling pathway and RET inhibitors in human cancer, Front Oncol, 12 (2022) 932353.

[9] J. Kim, D. Bradford, E. Larkins, L.H. Pai-Scherf, S. Chatterjee, P.S. Mishra-Kalyani, E. Wearne, W.S. Helms, A. Ayyoub, Y. Bi, J. Sun, R. Charlab, J. Liu, H. Zhao, D. Liang, S. Ghosh, R. Philip, R. Pazdur, M.R. Theoret, J.A. Beaver, H. Singh, FDA Approval Summary: Pralsetinib for the Treatment of Lung and Thyroid Cancers With RET Gene Mutations or Fusions, Clin Cancer Res, 27 (2021) 5452–5456.

[10] V. Subbiah, M.I. Hu, L.J. Wirth, M. Schuler, A.S. Mansfield, G. Curigliano, M.S. Brose, V.W. Zhu, S. Leboulleux, D.W. Bowles, C.S. Baik, D. Adkins, B. Keam, I. Matos, E. Garralda, J.F. Gainor, G. Lopes, C.C. Lin, Y. Godbert, D. Sarker, S.G. Miller, C. Clifford, H. Zhang, C.D. Turner, M.H. Taylor, Pralsetinib for patients with advanced or metastatic RET-altered thyroid cancer (ARROW): a multi-cohort, open-label, registrational, phase 1/2 study, Lancet Diabetes Endocrinol, 9 (2021) 491–501.

[11] J.F. Gainor, G. Curigliano, D.W. Kim, D.H. Lee, B. Besse, C.S. Baik, R.C. Doebele, P.A. Cassier, G. Lopes, D.S.W. Tan, E. Garralda, L.G. Paz-Ares, B.C. Cho, S.M. Gadgeel, M. Thomas, S.V. Liu, M.H. Taylor, A.S. Mansfield, V.W. Zhu, C. Clifford, H. Zhang, M. Palmer, J. Green, C.D. Turner, V. Subbiah, Pralsetinib for RET fusion-positive non-small-cell lung cancer (ARROW): a multi-cohort, open-label, phase 1/2 study, Lancet Oncol, 22 (2021) 959–969.

[12] V. Subbiah, P.A. Cassier, S. Siena, E. Garralda, L. Paz-Ares, P. Garrido, E. Nadal, J. Vuky, G. Lopes, G.P. Kalemkerian, D.W. Bowles, M. Seetharam, J. Chang, H. Zhang, J. Green, A. Zalutskaya, M. Schuler, Y. Fan, G. Curigliano, Pan-cancer efficacy of pralsetinib in patients with RET fusion-positive solid tumors from the phase 1/2 ARROW trial, Nat Med, 28 (2022) 1640–1645.

[13] F. Griesinger, G. Curigliano, M. Thomas, V. Subbiah, C.S. Baik, D.S.W. Tan, D.H. Lee, D. Misch, E. Garralda, D.W. Kim, A.J. van der Wekken, J.F. Gainor, L. Paz-Ares, S.V. Liu, G.P. Kalemkerian, Y. Houvras, D.W. Bowles, A.S. Mansfield, J.J. Lin, V. Smoljanovic, A. Rahman, S. Kong, A. Zalutskaya, M. Louie-Gao, A.L. Boral, J. Mazieres, Safety and efficacy of pralsetinib in RET fusion-positive non-small-cell lung cancer including as first-line therapy: update from the ARROW trial, Ann Oncol, 33 (2022) 1168–1178.

[14] D. Bradford, E. Larkins, S.L. Mushti, L. Rodriguez, A.M. Skinner, W.S. Helms, L.S.L. Price, J.F. Zirkelbach, Y. Li, J. Liu, R. Charlab, F.R. Turcu, D. Liang, S. Ghosh, D. Roscoe, R. Philip, A. Zack-Taylor, S. Tang, P.G. Kluetz, J.A. Beaver, R. Pazdur, M.R. Theoret, H. Singh, FDA Approval Summary: Selpercatinib for the Treatment of Lung and Thyroid Cancers with RET Gene Mutations or Fusions, Clin Cancer Res, 27 (2021) 2130–2135.

[15] V. Subbiah, J. Wolf, B. Konda, H. Kang, A. Spira, J. Weiss, M. Takeda, Y. Ohe, S. Khan, K. Ohashi, V. Soldatenkova, S. Szymczak, L. Sullivan, J. Wright, A. Drilon, Tumour-agnostic efficacy and safety of selpercatinib in patients with RET fusion-positive solid tumours other than lung or thyroid tumours (LIBRETTO-001): a phase 1/2, open-label, basket trial, Lancet Oncol, 23 (2022) 1261–1273.

[16] V. Subbiah, J.F. Gainor, G.R. Oxnard, D.S.W. Tan, D.H. Owen, B.C. Cho, H.H. Loong, C.E. McCoach, J. Weiss, Y.J. Kim, L. Bazhenova, K. Park, H. Daga, B. Besse, O. Gautschi, C. Rolfo, E.Y. Zhu, J.F. Kherani, X. Huang, S. Kang, A. Drilon, Intracranial Efficacy of Selpercatinib in RET Fusion-Positive Non-Small Cell Lung Cancers on the LIBRETTO-001 Trial, Clin Cancer Res, 27 (2021) 4160–4167.

[17] A. Drilon, G.R. Oxnard, D.S.W. Tan, H.H.F. Loong, M. Johnson, J. Gainor, C.E. McCoach, O. Gautschi, B. Besse, B.C. Cho, N. Peled, J. Weiss, Y.J. Kim, Y. Ohe, M. Nishio, K. Park, J. Patel, T. Seto, T. Sakamoto, E. Rosen, M.H. Shah, F. Barlesi, P.A. Cassier, L. Bazhenova, F. De Braud, E. Garralda, V. Velcheti, M. Satouchi, K. Ohashi, N.A. Pennell, K.L. Reckamp, G.K. Dy, J. Wolf, B. Solomon, G. Falchook, K. Ebata, M. Nguyen, B. Nair, E.Y. Zhu, L. Yang, X. Huang, E. Olek, S.M. Rothenberg, K. Goto, V. Subbiah, Efficacy of Selpercatinib in RET Fusion-Positive Non-Small-Cell Lung Cancer, N Engl J Med, 383 (2020) 813–824.

[18] A. Drilon, V. Subbiah, O. Gautschi, P. Tomasini, F. de Braud, B.J. Solomon, D. Shao-Weng Tan, G. Alonso, J. Wolf, K. Park, K. Goto, V. Soldatenkova, S. Szymczak, S.S. Barker, T. Puri, A. Bence Lin, H. Loong, B. Besse, Selpercatinib in Patients With RET Fusion-Positive Non-Small-Cell Lung Cancer: Updated Safety and Efficacy From the Registrational LIBRETTO-001 Phase I/II Trial, J Clin Oncol, 41 (2023) 385–394.

[19] L.J. Wirth, E. Sherman, B. Robinson, B. Solomon, H. Kang, J. Lorch, F. Worden, M. Brose, J. Patel, S. Leboulleux, Y. Godbert, F. Barlesi, J.C. Morris, T.K. Owonikoko, D.S.W. Tan, O. Gautschi, J. Weiss, C. de la Fouchardiere, M.E. Burkard, J. Laskin, M.H. Taylor, M. Kroiss, J. Medioni, J.W. Goldman, T.M. Bauer, B. Levy, V.W. Zhu, N. Lakhani, V. Moreno, K. Ebata, M. Nguyen, D. Heirich, E.Y. Zhu, X. Huang, L. Yang, J. Kherani, S.M. Rothenberg, A. Drilon, V. Subbiah, M.H. Shah, M.E. Cabanillas, Efficacy of Selpercatinib in RET-Altered Thyroid Cancers, N Engl J Med, 383 (2020) 825–835.

[20] S.Y. Wu, S. Sharma, K. Wu, A. Tyagi, D. Zhao, R.P. Deshpande, K. Watabe, Tamoxifen suppresses brain metastasis of estrogen receptor-deficient breast cancer by skewing microglia polarization and enhancing their immune functions, Breast Cancer Res, 23 (2021) 35.

[21] P.D. Bos, X.H. Zhang, C. Nadal, W. Shu, R.R. Gomis, D.X. Nguyen, A.J. Minn, M.J. van de Vijver, W.L. Gerald, J.A. Foekens, J. Massague, Genes that mediate breast cancer metastasis to the brain, Nature, 459 (2009) 1005–1009.

[22] D. Doheny, S. Sirkisoon, R.L. Carpenter, N.R. Aguayo, A.T. Regua, M. Anguelov, S.G. Manore, A. Arrigo, S.A. Jalboush, G.L. Wong, Y. Yu, C.J. Wagner, M. Chan, J. Ruiz, A. Thomas, R. Strowd, J. Lin, H.W. Lo, Combined inhibition of JAK2-STAT3 and SMO-GLI1/tGLI1 pathways suppresses breast cancer stem cells, tumor growth, and metastasis, Oncogene, 39 (2020) 6589–6605.

[23] A.T. Regua, S. Bindal, M.K. Najjar, C. Zhuang, M. Khan, A.B.J. Arrigo, A.O. Gonzalez, X.R. Zhang, J.J. Zhu, K. Watabe, H.W. Lo, Dual inhibition of the TrkA and JAK2 pathways using entrectinib and pacritinib suppresses the growth and metastasis of HER2-positive and triple-negative breast cancers, Cancer Lett, 597 (2024) 217023.

[24] S.R. Sirkisoon, R.L. Carpenter, T. Rimkus, D. Doheny, D. Zhu, N.R. Aguayo, F. Xing, M. Chan, J. Ruiz, L.J. Metheny-Barlow, R. Strowd, J. Lin, A.T. Regua, A. Arrigo, M. Anguelov, B. Pasche, W. Debinski, K. Watabe, H.W. Lo, TGLI1 transcription factor mediates breast cancer brain metastasis via activating metastasis-initiating cancer stem cells and astrocytes in the tumor microenvironment, Oncogene, 39 (2020) 64–78.

[25] D. Doheny, S. Manore, S.R. Sirkisoon, D. Zhu, N.R. Aguayo, A. Harrison, M. Najjar, M. Anguelov, A.O. Cox, C.M. Furdui, K. Watabe, T. Hollis, A. Thomas, R. Strowd, H.W. Lo, An FDA-Approved Antifungal, Ketoconazole, and Its Novel Derivative Suppress tGLI1-Mediated Breast Cancer Brain Metastasis by Inhibiting the DNA-Binding Activity of Brain Metastasis-Promoting Transcription Factor tGLI1, Cancers (Basel), 14 (2022).

[26] A. Gandillet, I. Vidal, E. Alexandre, M. Audet, M.P. Chenard-Neu, J. Stutzmann, B. Heyd, D. Jaeck, L. Richert, Experimental models of acute and chronic liver failure in nude mice to study hepatocyte transplantation, Cell Transplant, 14 (2005) 277–290.

[27] A. Liberzon, C. Birger, H. Thorvaldsdottir, M. Ghandi, J.P. Mesirov, P. Tamayo, The Molecular Signatures Database (MSigDB) hallmark gene set collection, Cell Syst, 1 (2015) 417–425.

[28] A. Liberzon, A. Subramanian, R. Pinchback, H. Thorvaldsdottir, P. Tamayo, J.P. Mesirov, Molecular signatures database (MSigDB) 3.0, Bioinformatics, 27 (2011) 1739–1740.

[29] L.J. Bendall, D.R. Green, Autopsy of a cell, Leukemia, 28 (2014) 1341–1343.

[30] S.H. Kaufmann, S. Desnoyers, Y. Ottaviano, N.E. Davidson, G.G. Poirier, Specific proteolytic cleavage of poly(ADP-ribose) polymerase: an early marker of chemotherapy-induced apoptosis, Cancer Res, 53 (1993) 3976–3985.

[31] Selpercatinib, LiverTox: Clinical and Research Information on Drug-Induced Liver Injury, Bethesda (MD), 2012.

[32] Pralsetinib, LiverTox: Clinical and Research Information on Drug-Induced Liver Injury, Bethesda (MD), 2012.

[33] P. Jagust, A.M. Powell, M. Ola, L. Watson, A. de Pablos-Aragoneses, P. Garcia-Gomez, R. Fallon, F. Bane, M. Heiland, G. Morris, B. Cavanagh, J. McGrath, D. Ottaviani, A. Hegarty, S. Cocchiglia, K.J. Sweeney, S. MacNally, F.M. Brett, J. Cryan, A. Beausang, P. Morris, M. Valiente, A.D.K. Hill, D. Vareslija, L.S. Young, RET overexpression leads to increased brain metastatic competency in luminal breast cancer, J Natl Cancer Inst, (2024).

[34] D. Vareslija, N. Priedigkeit, A. Fagan, S. Purcell, N. Cosgrove, P.J. O’Halloran, E. Ward, S. Cocchiglia, R. Hartmaier, C.A. Castro, L. Zhu, G.C. Tseng, P.C. Lucas, S.L. Puhalla, A.M. Brufsky, R.L. Hamilton, A. Mathew, J.P. Leone, A. Basudan, L. Hudson, R. Dwyer, S. Das, D.P. O’Connor, P.G. Buckley, M. Farrell, A.D.K. Hill, S. Oesterreich, A.V. Lee, L.S. Young, Transcriptome Characterization of Matched Primary Breast and Brain Metastatic Tumors to Detect Novel Actionable Targets, J Natl Cancer Inst, 111 (2019) 388–398.

[35] E. Tabouret, P. Metellus, A. Goncalves, B. Esterni, E. Charaffe-Jauffret, P. Viens, A. Tallet, Assessment of prognostic scores in brain metastases from breast cancer, Neuro Oncol, 16 (2014) 421–428.

