## Supplemental Figures and Table for "RET Receptor Tyrosine Kinase Promotes Breast Cancer Metastasis to the Brain and RET Inhibitors Pralsetinib and Selpercatinib Suppress Breast Cancer Brain Metastases"

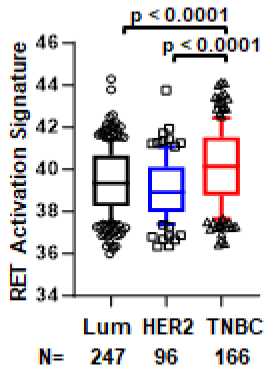


**Supplemental Figure 1: RET activation signature is significantly higher in patients with TNBC subtype.** To further investigate the clinical relevance of these observations, we conducted data mining analysis using our curated GEO breast cancer patient's dataset and analyzed RET activation across breast cancer subtypes.

| | | IC50 (μM) | |
|---|---|---|---|
| Subtype | Cell line | Pralsetinib | Selpercatinib |
| HER2-enriched | SKBR3 | 9.63 | 27.48 |
| | SKBRM | 9.97 | 25.80 |
| TNBC | MDA-231 | 19.79 | 20.8 |
| | MDA-231-BrM | 12.09 | 29.53 |
| | CN34 | 10.80 | 19.87 |
| | CN34-BrM | 18.04 | 28.07 |

**Supplemental Figure 2: Breast cancer cells and their brain-tropic variants are sensitive to RET inhibitors (RETi).** Representative IC50 values of RETi derived from treatment of HER2-enriched parental SKBR3 cell line and its brain-tropic variant SKBRM, and the TNBC cell lines MDA-MB-231 or CN34 and their brain-tropic variants MDA-MB-231-BrM or CN34-BrM, respectively.
